# DDX3X acts as a selective dual switch regulator of mRNA translation in acute ER stress

**DOI:** 10.1101/2025.06.08.658532

**Authors:** Abd-El Monsif A. Shawky, Allison Scarboro, Josef Mick, Mahmoud Dondeti, Kenneth Avanzino, Constantine A. Simintiras, Maria Hatzoglou, Anastasios Vourekas

## Abstract

In eukaryotes, regulation of mRNA translation initiation greatly impacts gene expression, and is critical for cellular stress responses. DDX3X is a ubiquitous DEAD-box RNA helicase whose precise role in 5‘ UTR scanning and start codon decoding in non-stressed and stressed cells is still elusive. Here we show that DDX3X engages with thousands of mRNAs as part of the eIF4F-mediated 48S scanning complex, simultaneously acting to promote or suppress translation of select mRNAs in non-stressed conditions, and switches this regulation in opposite directions in acute ER stress. We find distinct DDX3X binding patterns of differentially regulated mRNAs, which lead us to identify N4-acetylation of cytidines surrounding the start codon as an accompanying feature of mRNAs subject to DDX3X-mediated selective dual regulation. Our findings illuminate the role of DDX3X in stress response and highlight a novel connection between an RNA helicase and a post-transcriptional modification in regulating mRNA translation.

## Introduction

mRNA translation plays a central role in the regulation of gene expression, with much of this control occurring at the level of initiation. Translation initiation requires the scanning of the mRNA 5‘ <u>u</u>ntranslated region (UTR) for the identification of the start codon by the 48S scanning complex, which forms when the 43S pre-initiation complex interacts with the eIF4F-activated mRNA ^1,2^. Translocation of the 48S complex and cognate and near-cognate start codon recognition is influenced by local mRNA secondary structure, which, depending on its positioning, may positively or negatively affect translation initiation, and the selection of the translation initiation site ^2,3^.

DEAD-box RNA helicases^4^ play critical roles in this process, by resolving secondary structures, and ensuring directionality of scanning: eIF4A (an eIF4F component) is a general purpose helicase required for translation of most mRNAs ^5–9^, while the mammalian DDX3X and its yeast counterpart Ded1p target select mRNAs to promote mRNA translation ^10–17^. Elucidating DDX3X mRNA binding and selectivity in translation regulation is important for understanding stress response and disease pathogenesis, as DDX3X has been implicated in both.

Biochemical ^10,18–22^ and structural studies have established a framework for understanding the molecular and biochemical properties of DDX3X as an RNA helicase, primarily for ATP-dependent unwinding of RNA secondary structure; molecular structures have revealed contacts of the conserved helicase core only with the substrate RNA backbone and 2‘-OH and no base-specific contacts ^21,23–25^ suggesting that DDX3X helicase core does not have an inherent sequence preference for its mRNA targets. This finding is corroborated by DDX3X Crosslinking immunoprecipitation and deep sequencing (CLIP-Seq) studies, which thus far have not identified enriched motifs within DDX3X bound mRNA sequences ^14–16^.

Cellular stress activates the Integrated Stress Response (ISR) leading to phosphorylation of eIF2α ^26^, reduced ternary complex (TC) levels and global mRNA translation suppression during the initial acute phase ^27^. This translational suppression is believed to promote the hallmark formation of cytoplasmic membraneless organelles known as stress granules, where DDX3X^16,28^ is localized along with stalled ribosomes and translation initiation factors^29^. Concurrently and despite low TC levels, translation of certain mRNAs is stimulated, exemplified by mammalian ATF4, a master transcription regulator of ISR ^30,31^. Notably, during the acute phase of endoplasmic reticulum (ER) stress, translation of ATF4 and other upstream ORF containing mRNAs is DDX3X-dependent^32,33^. Moreover, DDX3X increases cell viability and recovery after oxidative stress^34^. Taken together, the presence of DDX3X in stress granules, and ATF4 translation stimulation by DDX3X in the acute phase of ER stress, suggest a critical dual role for DDX3X in stress response to support cell survival. In line with this notion, stress-induced acetylation of DDX3X moderates its recruitment in stress granules^28^. However, the specific translational changes induced by DDX3X that underlie this pro-survival function remain to be fully elucidated. Additionally, recent studies have uncovered multilayered translational control of stress response that engages eIF4F-dependent and independent pathways. These mechanisms operate not only temporally, separating the acute from the chronic phase of ER stress^35^, but also across distinct branches of the ISR: the canonical ISR characterized by increased eIF2α-P and low eIF2B activity, and a “split” ISR, distinguished by low eIF2B activity, as observed in pathological conditions such as vanishing white matter disease^36,37^. Understanding DDX3X’s role within this regulatory landscape will offer new insights into the coordination and selectivity of stress-responsive translation.

DDX3X mutations have been implicated in cancerogenesis and neurodevelopmental disorders like DDX3X syndrome^38^. Cell and animal models have shown variable effects of DDX3X mutations, which impact phenotype, mRNA abundance and translation, and stress granule formation ^14,16,39–41^. Evidence points to altered stress responses due to DDX3X mutations, which contributes to cancerogenesis, albeit through different routes. For example, loss of function of DDX3X in medulloblastoma is thought to result in reduced sensing of oncogenic stress that promotes tumor formation^41^. In contrast, an initial loss of DDX3X followed by a subsequent ectopic expression of the Y-linked DDX3X paralogue DDX3Y is thought to buffer proteotoxic stress in MYC-driven lymphomas and promote tumor formation^42^. Therefore, understanding the role of DDX3X in stress response can contribute to an improved understanding of its role in cancer pathogenesis.

Several studies have investigated mRNA features that influence DDX3X-mediated regulation. In addition to the absence of enriched sequence motifs in DDX3X-bound regions^14–16^, increased length and GC content of 5‘ UTR have been reported as common features of many mRNAs that depend on DDX3X for their translation^15,17,43^. However, these features have limited predictive value. Ribosomal protein mRNAs are DDX3X-dependent mRNAs and harbor short 5‘

UTRs with inhibitory structures^44^. Additionally, the 5‘ UTR alone may not be sufficient to confer DDX3X sensitivity in reporter assays^43^, suggesting that additional contextual factors may be important.

Recent data suggest that DDX3X, Ded1p and plant orthologs resolve structured elements downstream of alternative translation initiation sites, impacting main ORF translation and stress response ^3,13,45^. Moreover, post-transcriptional modifications of mRNA (PTMs) like N^4^-acetylcytidine (ac4C) have been shown to regulate mRNA translation and stability, with the position of the acetylated residues relative to the start codon playing and important role ^46,47^. The potential interplay between transient PTMs and dynamic mRNA secondary structure could provide the basis for understanding how RNA helicases such as DDX3X mediate selective translation regulation under changing cellular conditions.

Here we use endogenous DDX3X CLIP-Seq and DDX3X knockdown coupled with polysome profiling to identify mRNAs regulated by DDX3X under basal and ER stress conditions. We find that DDX3X acts as a dual switch regulator by controlling suppression and activation of mRNA translation to serve the needs of non-stressed cells, but also to prepare and execute their response to acute ER stress. We identify distinct cytidine acetylation patterns around start codons of mRNAs that are bidirectionally regulated by DDX3X, and uncover that mRNAs suppressed by DDX3X in non-stressed cells also require acetylation for their suppression, indicating a functional significance of ac4C patterns as a possible dynamic mRNA feature assisting DDX3X translational regulation.

## Materials and methods

### Cell culture

Mouse embryonic fibroblasts (MEFs) were grown in high glucose Dulbecco’s modified Eagle’s medium (DMEM) supplemented with 10% defined fetal bovine serum (FBS) (Gibco), 2 mM L-glutamine, 100 units/mL penicillin, and 100 μg/mL streptomycin.

293FT cells (Thermo R70007) and 293T cells (ATCC CRL-3216) were grown in high glucose Dulbecco’s modified Eagle’s medium (DMEM, Thermo 11965084) supplemented with 10% FBS (Atlas Biologicals, F-0500-DR), 1x Glutamax (Thermo), MEM non-essential amino acids 1X (Thermo 11140050) and Sodium Pyruvate (Thermo 11360070) 1 mM final concentration. Cells were grown at 37° C under 5% CO2. The cells were not allowed to achieve full confluency before they were split.

Acute ER stress was induced by treating cells with full media containing 400 nM Thapsigargin (Tg, Sigma T9033) for 1 or 1.5 hours, as noted; control treatment was with DMSO alone.

### Chemicals, reagents, antibodies

RNase H (Promega, M428A), rRNase inhibitor (Promega, N2515), RNase T1 (Thermo, EN0542), RQ1 DNase (Promega, M610A), Sodium Dedocyl Sulfate (SDS) (Fisher, BP1311-200 mL), 4X NUPAGE SDS loading buffer (Invitrogen, NP0007) supplemented with 1/10 v/v β-mercaptoethanol, Sodium Acetate pH(5.2) (Invitrogen, AM9740), Sodium Chloride (Fisher, BP358-1Kg), Sodium deoxycholate (EMD Millipore, 264103-25g), Sodium Pyruvate 100 mM (Thermo, 11360070), D-Sucrose (Fisher, BP220-1 Kg), SuperScript IV Reverse Transcriptase (Invitrogen, 18080093), SYBR green Mix (Applied biosystems, A46109), T4 Polynucleotide Kinase (T4 PNK; NEB) (provider., cat #), T4 RNA ligase (Thermo EL0021), Thapsigargin (Sigma T9033), Triton X-100 (Fisher, BP151-500mL)EDTA ultrapure (Invitrogen, 15575020), Dithiothreitol (DTT) (100 mM) (Roche, 10278200), AccuPrime^TM^ Pfx SuperMix (Invitrogen, 12344-040), Cycloheximide (MP Biomedicals, 194527), Antarctic phosphatase (New England Biolabs, M0289L), Anti-eIF2αAntibody (Cell Signaling technology, 5324S), Rabbit polyclonal anti-DDX3X (custom antibody, Genscript), anti-tubulin (Cell Signaling technology, 3837S), anti-ATF4 (Proteintech., 10835-1-AP), anti-NAT10 (Proteintech 13365-1-AP), anti-eIF2αP Ser-52 (Cell Signaling technology, 3398S), were used at 1:1000 dilution. Goat anti-rabbit StarBright blue 700 (Bio-Rad 12004162), IRDye® 800CW Goat anti-Mouse (Li-Cor 926-32210) were used at 1:5000.

### DDX3X, NAT10 and control knockdown

293FT cells were seeded at ∼ 30% confluency and transfected with DDX3X dsiRNA (IDT hs.Ri.DDX3X.13.3), NAT10 siRNA (Dharmacon, ON-TARGET plus Human NAT10 (55226) siRNA - SMARTpool) or non-targeting control dsiRNA (IDT, NC1) at a final concentration of 30 nM. The oligo was mixed with RNAiMax Lipofectamine (Thermo 13778075) or DharmaFECT1 (Horizon Discovery Biosciences T-2001-03) and optiMEM (Thermo 31985062) as per manufacturer’s instructions, and the transfection mixture was added to the full medium and transferred to cell culture plates in which the appropriate dilution of cell suspension was afterwards added. After 48 hours, cells were washed with ice cold PBS, and were harvested by scraping, followed by washing with PBS and centrifugation at 4,500 RPM for 5 min at 4° C. DDX3X and NAT10 knockdown (≥65%) was confirmed via western blotting.

### Polysome profiling and fractionation

We prepared 10%-50% sucrose gradients in polyallomer tubes for SW-41 rotor using the Gradient Master 108, and collected the polysome profiles using the Piston Fractionator with a TRIAX Flowcell (260 nm) (BioComp). 293FT cells were seeded in 100-mm culture dishes (3 dishes per polysome profile sample) and grown to 80% confluency. 1.5 h prior to collection, cell culture media was replaced with full media containing DMSO (non-stressed condition) or 400 nM Thapsigargin (acute ER stress). Cells were incubated for the last 5 min of the treatment with cycloheximide (CHX) at a final concentration of 100 μg/mL. Cells were then washed twice with ice cold phosphate-buffered saline supplemented with CHX and centrifuged at 4,500 X g for 5 min at 4° C. Cell pellets were lysed in 800 μL of Hypotonic buffer (5 mM Tris HCl pH 7.5, 2.5 mM MgCl_2_, 1.5 mM KCl, 0.5% Triton X, 0.5% sodium deoxycholate, 100 μg/mL CHX, 2 mM DTT, 200 units/mL RNase inhibitor, 1X protease inhibitor cocktail). Cells were resuspended in the hypotonic buffer either by pipetting or vortexing for 5 sec followed by centrifugation at 17,000 xg for 7 min at 4° C. The supernatant (post-nuclear lysate) OD was measured at 260 nm to determine the amount of lysate needed to be added to sucrose gradient. Samples of the post-nuclear lysates were also kept for western blots and input RNA extraction 500 mL from the top of the sucrose gradient was removed and the lysate volume per condition was adjusted so they contain the same OD between 10 to 20 at 260 nm and added to the top of the sucrose gradient. Each gradient was weighed and balanced before ultracentrifugation at 222,222 xg for 2 hours at 4° C while using low brake option. 14 fractions of 800 μL were collected.

For experiments involving acquisition of monosomes, cells were treated with a final concentration of 2µg/mL harringtonine (HRN) for 10 min prior to the 5-min treatment with cycloheximide. The HRN treatment began 15 minutes prior to the end of the 1.5h stress (thapsigargin) or rest (DMSO) treatment. Fraction collection was adjusted so that the entire monosomal peak was collected in two fractions. RNA extraction from monosomes and input RNA was performed as described below.

### RNA extraction from polysome profiles, reverse transcription, and qPCR

Immediately after polysome fractions were collected, 400 μL of each fraction was mixed with equal volume TRIZOL LS (Thermo) and kept at-80 °C. RNA extraction was carried out by manufacturer’s instructions. RNA was treated with RQ1 DNase (Promega), followed by phenol extraction and ethanol precipitation. The concentration and the purity of RNA was measured using nanodrop and pure RNA with a 260/280 ratio within 1.8 - 2.0 was acquired. Equal volume of RNA samples were spiked with 500 ng of Renilla luciferase mRNA (transcribed *in vitro* from pRL Renilla Luciferase encoding plasmid, Promega), in place of an endogenous reference mRNA, for calculating relative mRNA quantities.

Spiked RNA from polysome fractions were reverse transcribed using random hexamers and Superscript III for 10 min at 25° C, for 50 min at 50° C then finally for 5 min at 85° C. 1 μL of RNase H was added then the mixture was incubated for 20 min at 37° C. The cDNA was used for qPCR with the PowerTrack SYBR Green Master Mix (Thermo), on a QuantStudio3 (Thermo). mRNA levels were calculated with the ΔΔCt method using QuantStudio™ Design & Analysis Software.

For RT-qPCR experiments following Harringtonine treatment specifically, the monosome peak was collected in two fractions and 200 μL of the mixed fractions was used for RNA extraction using Trizol LS. In parallel, 50 μL of the post nuclear lysate used for the gradient density centrifugation was used for RNA extraction (input). 10 ng of RNA extracted from the monosome peak and the input sample (spiked with Renilla luciferase mRNA) were used for cDNA synthesis using qScript Ultra SuperMix (Quantabio, 95217-100). 0.25 μL of the cDNA reaction was used as a template per qPCR sample.

### DDX3X immunoprecipitation and peptide elution for mass spectrometry

8E5 cells were lysed by sonication on ice in RSB150 Lysis and IP buffer (9 μL of buffer per 1 mg of cells): 50 mM Tris pH 7.5, 150 mM NaCl, 0.5% NP-40 IGEPAL, 0.1% Triton X-100, 1.25 mM EDTA, with Halt™ Protease and Phosphatase Inhibitors (Thermo 78442), and 0.4U/μL RNasin. The lysate was centrifuged at 16000xg for 20 min. 800 μL of cleared lysate was mixed with of 80 μL of Protein A Dynabeads slurry (Thermo 10002D) bound with 6 μg of DDX3X Ab, and incubated for 2 hours at 4°C. Subsequently, for the RNase treatment sample, the IP beads were washed once with RSB150 and then incubated with 1 μL undiluted RNase T1 (1000U) and incubated for 10 min at RT. IP beads were then washed 3 times with RSB150, and eluted for 25 min at RT by light shaking with RSB750 (same composition as RSB150 but with 750 mM of NaCl) plus 2.5 μg/μL of the DDX3X antigenic peptide used to raise the Ab in rabbits (RYIPPHLRNREATK). Standard elution is performed with reducing 4X LDS sample buffer (Thermo) at 70 °C for 12 min. To submit a sample for mass spectrometry, 30 μL of the elution sample was run for 9 min on a 4-12% NuPage gel (Thermo) until the entire volume is ion the gel, and then a thin slice of the gel was cut and submitted to Taplin Mass Spectrometry facility at Harvard.

### Western blotting

Cells were harvested by trypsinization or scraping and washed with ice cold PBS before lysis. Cell lysates for western blot were prepared as described above for IP. Lysates were mixed with equal volume of reducing 4X LDS sample buffer (Thermo) at 70 °C for 12 min. Samples were run on 4-12% Bis-Tris NuPage gels (Thermo) and transferred on nitrocellulose (Cytiva, 10600001). Single or dual fluorescence detection of protein bands was performed on a ChemiDoc MP system (Bio-Rad). The intensity of the immunoreactive bands was determined using image lab software for densitometric analysis.

### CLIP-Seq

MEFs and 293T cells were grown to ∼80% confluency and treated with control (DMSO) or Thapsigargin (400 nM) containing full medium to induce acute ER stress. The cells where then collected by trypsinization and kept in ice-cold HBSS during UV crosslinking (254 nm, 3×400 mJ/cm^2^). To ensure that the procedure is not inducing stress in the control cells, we checked levels of eIF2α-P and ATF4 by western blot, and performed DDX3X immunofluorescence to observe possible induction of stress granules (Figure S1). For the IP step our own custom rabbit polyclonal DDX3X antibody was used. The rest of the CLIP-Seq procedure was performed as previously described ^48^.

### RNA-seq library preparation

To prepare polyA mRNA RNA-Seq libraries, approximately 800 ng of high-quality total RNA was used per sample. The quality was determined with a Nanodrop spectrophotometer, and only RNA that had a OD260/280 of 1.8-2.0 was used. We followed the Illumina Stranded mRNA Prep, Ligation Reference Guide, Document # 1000000124518 V03 protocol without changes.

### Cell viability assay

At day 1 cells were treated with RNAi (30 nmol of oligo) as described above, and after 48h cells were washed and treated with CPA (200 μM) or DMSO (negative control) in full medium for 1.5h. The cells were then washed with full medium, trypsinized, counted and plated on 96 well plates in sequential dilutions in three replicates. Cell viability was assessed at 24h and 48h using CellTiter-Glo Luminescent Cell Viability Assay (Promega, G7570), using a Veritas Microplate Luminometer. DDX3X and control mock KD were assessed at 24h and 48h with parallel cell samples.

### Bioinformatic analysis

CLIP-Seq libraries were demultiplexed using Cutadapt v. 4.1 with python 3.9.13. Fixed adapter sequences at the 5‘ ends of reads (*<u>AGGGAGGACGATGCGG</u>*) were trimmed using Cutadapt v. 4.1 with python 3.9.13. Options-j 7-n 1-e 0.15 accelerated the trimming procedure and allowed for one adaptor at a time to be trimmed while limiting to 2 mismatches withing each sequence. Next, fixed adapter sequences at the 3‘ ends of reads (*<u>GTGTCAGTCACTTC</u>*) were removed using the same settings.

Thereafter reads were collapsed to remove PCR amplified reads and obtain non-redundant reads using Seqcluster v. 1.2.9. Unique molecular identifiers (UMIs) were removed using Umi tools v. 1.1.2 and selecting the option to extract NNNNG patterns during UMI removal. Filtered reads were processed using CLIPSEQTOOLS with the preprocess option to only retain reads with a minimum length of 16 nucleotides while limiting to 1 mismatch per adaptor (this extra filtering step removes tandem adaptor sequences that can be left over from previous trimming steps). Then, read alignment was performed using STAR aligner v. 2.7.0d with the runThreadN 10 option to use 10 cores. A.sam file with all attributes was created using the “outSAMattributes All” option, and allowing matches for reads that are at least 15 nucleotides with outFilterMatchNmin 15, allowing only 90% matching with outFilterMatchNminOverLread 0.9 and obtaining a second sam file containing the Transcriptome results (mapped to the transcriptome) using the option quantMode TranscriptomeSAM. All libraries were aligned to both genomes hg19 and hg38 after creating indices for both genomes. STAR aligned reads (sam files) were sorted and only one single record was kept for multimappers using clipseqtools preprocess with the option cleanup_alignment. Then, filtered STAR aligned reads were loaded with the alignments in an SQLite database using clipseqtools preprocess with the option sam_to_seqlite. The newly created database was annotated permitting deletions using the option annotate_with_deletions. Furthermore, the reads were annotated for positions on genome using UCSC gene parts GTF (Gene Transfer Format) file using the option annotate_with_genic_elements. Annotation of alignments contained within regions from a BED/SAM file from rmsk.bed file was obtained using the option annotate_with_file. We used the option count_reads_on_genic_elements to map the reads and count them on transcripts, genes, exons, and introns, while we measured the distribution of reads along the length of the 5‘UTR, CDS, and 3‘UTR using the option distribution_on_genic_elements. We counted the number of reads on genes, repeats, exons, introns, 5‘UTR, CDS, 3‘UTR using the option genomic_distribution. Python custom scripts used in this study to create the meta-mRNA deep sequencing coverage plots, the log2 fold change scatterplots, and others can be found in the following GitHub page: https://github.com/Abdelmonsif/DDX3X.

Gene counts tables were used to calculate TPM (transcripts per million reads). For the polysome analysis from 293FT cells, we only kept transcripts that had a TPM value of 5 or above in any one of our polysome profiling libraries and had a length of 350 bases or longer. We calculated the fold change for each treatment using the log2 of the stress condition against the log2 of the control condition or the log2 of the gene knockdown experiment against the log2 of the mock knockdown experiment. To avoid the redundancy of the transcripts in the same gene, we chose the longest transcript that had the highest expression profile to represent each gene.

## Results

### Pervasive and distinct DDX3X binding of mRNAs translationally regulated upon acute ER stress

To capture endogenous DDX3X without transgene expression, we developed a custom rabbit polyclonal antibody against DDX3X, which efficiently and specifically depletes the protein from mouse (and human) cell lysates (Figures 1A and S1A). To identify bona fide mRNA targets of endogenous DDX3X, we performed CLIP-Seq ^48^ in mouse embryonic fibroblasts (information about high throughput sequencing libraries generated in this study can be found in Table S1), after control treatment (DMSO) and 1 h treatment with thapsigargin (Tg), which induces acute ER stress and eIF4E (eIF4F) dependent translational changes^35^ (Figure 1B). Because UV exposure can also induce stress, we verified by western blot (WB) that in control crosslinked cells, although eIF2α phosphorylation is mildly increased, this is not accompanied by ATF4 protein synthesis ^49^, while Tg treated cells show robust eIF2α phosphorylation and ATF4 synthesis (Figure 1B). Moreover, UV crosslinking as part of the CLIP protocol does not induce accumulation of DDX3X in stress granules (Figures S1B-S1D). The above support the applicability of CLIP-Seq for capturing mRNA translation regulation by DDX3X.

**Figure 1.**
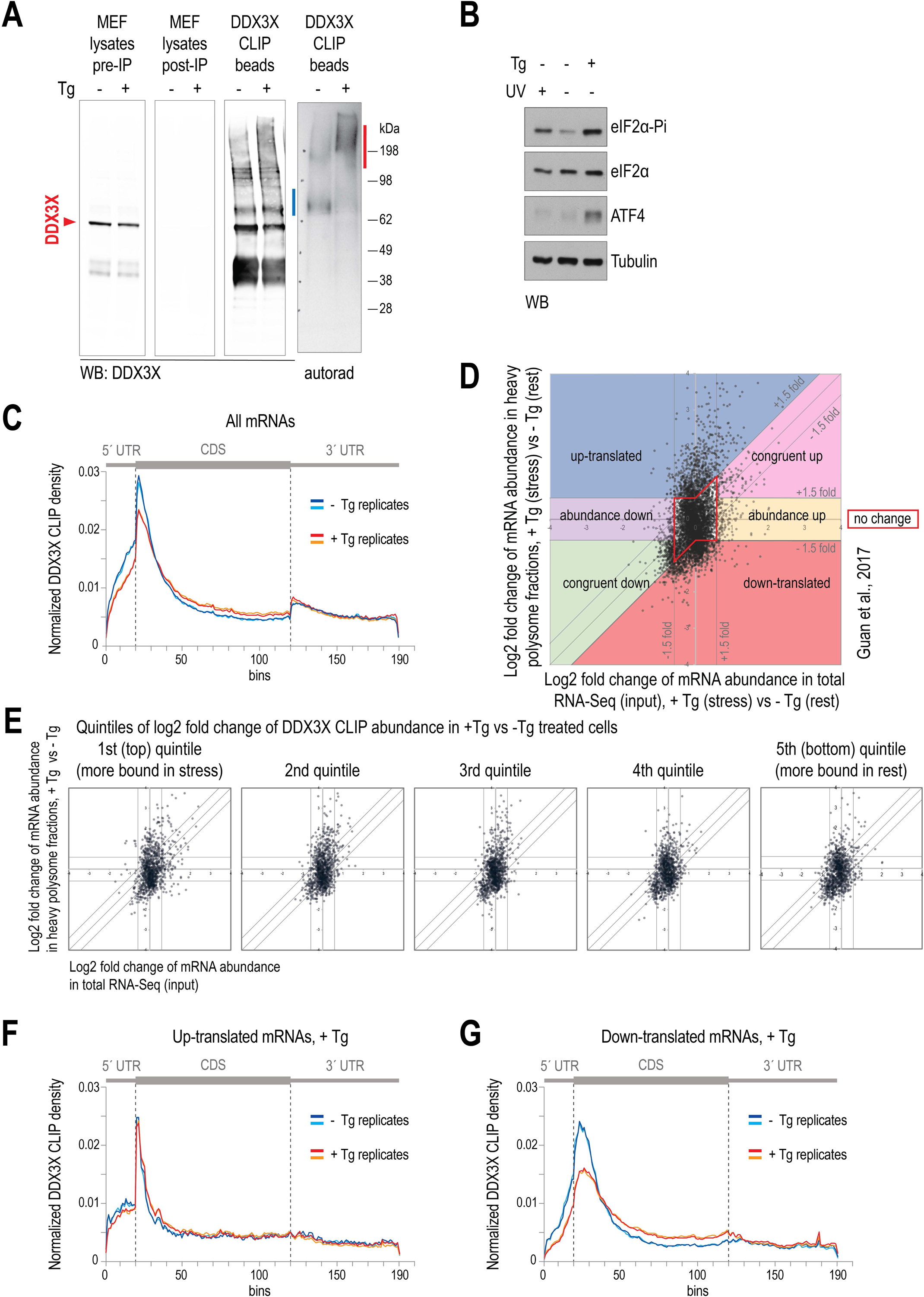
DDX3X CLIP-Seq in MEFs, mRNA regulation and DDX3X CLIP density on the meta-mRNA in ER stress (A) Western blot of MEF lysates and IP beads eluates for DDX3X CLIP, and autoradiography of IP beads eluates to visualize crosslinked DDX3X-RNA complexes (highlighted in blue for-Tg and red for +Tg). (B) Western blot for indicated proteins in MEF lysates used for DDX3X CLIP. (C) DDX3X CLIP density across the meta mRNA, for all mRNAs in-Tg and +Tg treated cells. (D) Polysome profiling mRNA scatterplot for changes in abundance and polysome association in MEFs treated with Tg compared to no Tg treatment. The 7 highlighted territories demarcate mRNAs with changes in heavy polysome association and/or mRNA abundance, or no change thereof. Data from Guan et al, 2017. (E) Polysome profiling mRNA scatterplot as in (D) but plotted separately for the quintiles of DDX3X-bound mRNAs ranked by the log2 fold change in DDX3X CLIP abundance in +Tg treated cells vs-Tg cells. (F), (G) DDX3X CLIP density across the meta mRNA, for up-translated mRNAs (F), and down-translated mRNAs (G), in-Tg and +Tg treated cells, as the mRNAs are demarcated in (D).

We detected strong radioactive signals from crosslinked RNA-DDX3X complexes in both control and stressed cells (Figures 1A and S1A), indicating that DDX3X does not globally dissociate from its mRNA targets under stress. Notably, Tg treatment caused a shift of the radioactive signal toward higher molecular weights (Figure 1A), underscoring the importance of autoradiography for optimal acquisition of protein-bound RNA in different conditions. RNA was extracted from the major radioactive bands (Figure 1A), and two biological replicates were generated per condition. The majority of CLIP-Seq reads (∼68% on average) mapped to mRNA exons (Figure S2A), consistent with the predominantly cytoplasmic localization of DDX3X observed by immunofluorescence (Figures S1B–S1D). We plotted CLIP-Seq read density across meta-mRNAs (Figure 1C), using binning based on the median nucleotide length of each mRNA region (5′ UTR, CDS, 3′ UTR) to maintain a uniform 10-nt resolution across segments (see Methods). Replicates showed high concordance and revealed distinct differences between conditions (Figure 1C). In control cells, DDX3X CLIP density gradually increases along the 5′ UTR and peaks in the early CDS, suggesting that binding at and just downstream of the start codon may be functionally as important as binding within the 5′ UTR. Upon Tg-induced stress, we observed a marked reduction in DDX3X CLIP density at both the 5′ UTR and the start codon region, accompanied by a modest increase in CDS binding. These changes suggest that DDX3X is involved in 5′ UTR scanning and start codon recognition, and that its binding pattern reflects the regulation of these processes during ER stress.

We proceeded with the simple hypothesis that DDX3X has one prominent contribution in mRNA translation regulation in stress, manifested by increased mRNA binding to promote translation and/or reduced mRNA association (possibly by DDX3X sequestration in stress granules ^50^) which results in down regulation of translation of DDX3X-dependent mRNAs. To test that, we sorted the DDX3X-bound mRNAs (CLIP-Seq abundance > 1 TPM in any library) by log2 fold change of DDX3X CLIP abundance in +Tg vs-Tg, and split the list into quintiles. We then plotted our previously published polysome profiling data from Tg treated MEFs^35^ (Figure 1D) separately for the quintiles of DDX3X-bound mRNAs (most to least abundance in +Tg vs-Tg treated cells), which revealed that, contrary to our hypothesis, change in DDX3X CLIP abundance in +Tg vs-Tg cells (Figure 1E), or absolute DDX3X CLIP abundance (Figures S2B and S2C) is not notably associated with up-translated or down-translated mRNAs, or any other mRNA regulation category (congruent-up, congruent-down, abundance-up, abundance-down, no change in +Tg vs-Tg cells). Importantly, mRNAs whose translation does not change in stress are also bound by DDX3X (area outlined in red in Figure 1D). The above suggest that a simple model in which DDX3X is either recruited to, or stripped from mRNAs to promote or to suppress mRNA translation is insufficient to explain the mechanism of regulation of this helicase in MEFs under acute ER stress.

We asked whether DDX3X distribution on the meta-mRNA could provide insights into its role in translational regulation during ER stress. To investigate this, we plotted CLIP density separately for mRNAs that are up-or down-translated in Tg-treated cells (Figures 1F and 1G). While overall DDX3X binding remains enriched around the start codon, distinct patterns emerge. Notably, up-translated mRNAs under stress diverge from the aggregate meta-mRNA profile (Figure 1C), showing relatively uniform DDX3X binding across the 5′ UTR, followed by an abrupt ∼3-fold increase precisely at the start codon and downstream bins. This sharp peak is followed by an equally steep decline, reaching the lowest DDX3X levels within the first fifth of the CDS and remaining low throughout the remaining mRNA (Figure 1F). This pattern remains unchanged in Tg-treated cells, and supports a functional role for DDX3X not only in the 5′ UTR but also at and immediately downstream of the start codon in promoting translation. In contrast, down-translated mRNAs display a gradual increase in DDX3X binding across the 5′ UTR, peaking shortly after the start codon and declining after the tenth CDS bin. Under stress, these transcripts show marked depletion of DDX3X in the 5′ UTR and around the start codon, consistent with reduced translation initiation (Figure 1G)

DDX3X binding is also enriched at and downstream of the start codon, with distinct pattern changes under Tg-induced stress observed in mRNAs showing no change in abundance or translation, as well as in those with congruent changes in both (Figures S3A–S3E). However, changes in mRNA abundance likely involve additional or secondary mechanisms beyond scanning and initiation. Therefore, we focus our subsequent analysis on transcripts regulated solely at the level of translation—the predominant mode of regulation during acute ER stress—as reflected by widespread changes in polysome association (y-axis), with relatively modest and fewer changes in abundance (Figure 1D).

Collectively, our results indicate that DDX3X predominantly binds mRNAs in the context of 5′ UTR scanning and translation initiation. CLIP meta-mRNA density plots implicate the 5′ UTR, start codon, and immediately downstream region as key sites where DDX3X exerts its regulatory role during acute ER stress. Notably, DDX3X binding at the start codon remains elevated in up-translated mRNAs but decreases in down-translated mRNAs under stress, suggesting that stress-induced changes in DDX3X binding patterns are closely tied to translational outcomes. This raises the question of how DDX3X actively contributes to both positive and negative regulation of translation. Altogether, our findings support a model in which DDX3X functions primarily through its role in the 48S scanning complex, rather than through changes in global mRNA binding levels.

### Bidirectional control of mRNA translation by DDX3X revealed by knockdown and polysome profiling

To explore the conservation of DDX3X regulatory mechanisms, we performed endogenous DDX3X CLIP-Seq in human 293T cells (Figure S4A). Genomic distribution of CLIP reads confirmed predominant binding to processed mRNAs (Figure S4B). The meta-mRNA CLIP density closely resembled that in MEFs, with increasing signal across the 5′ UTR and a peak just downstream of the start codon (Figure 2A).

**Figure 2.**
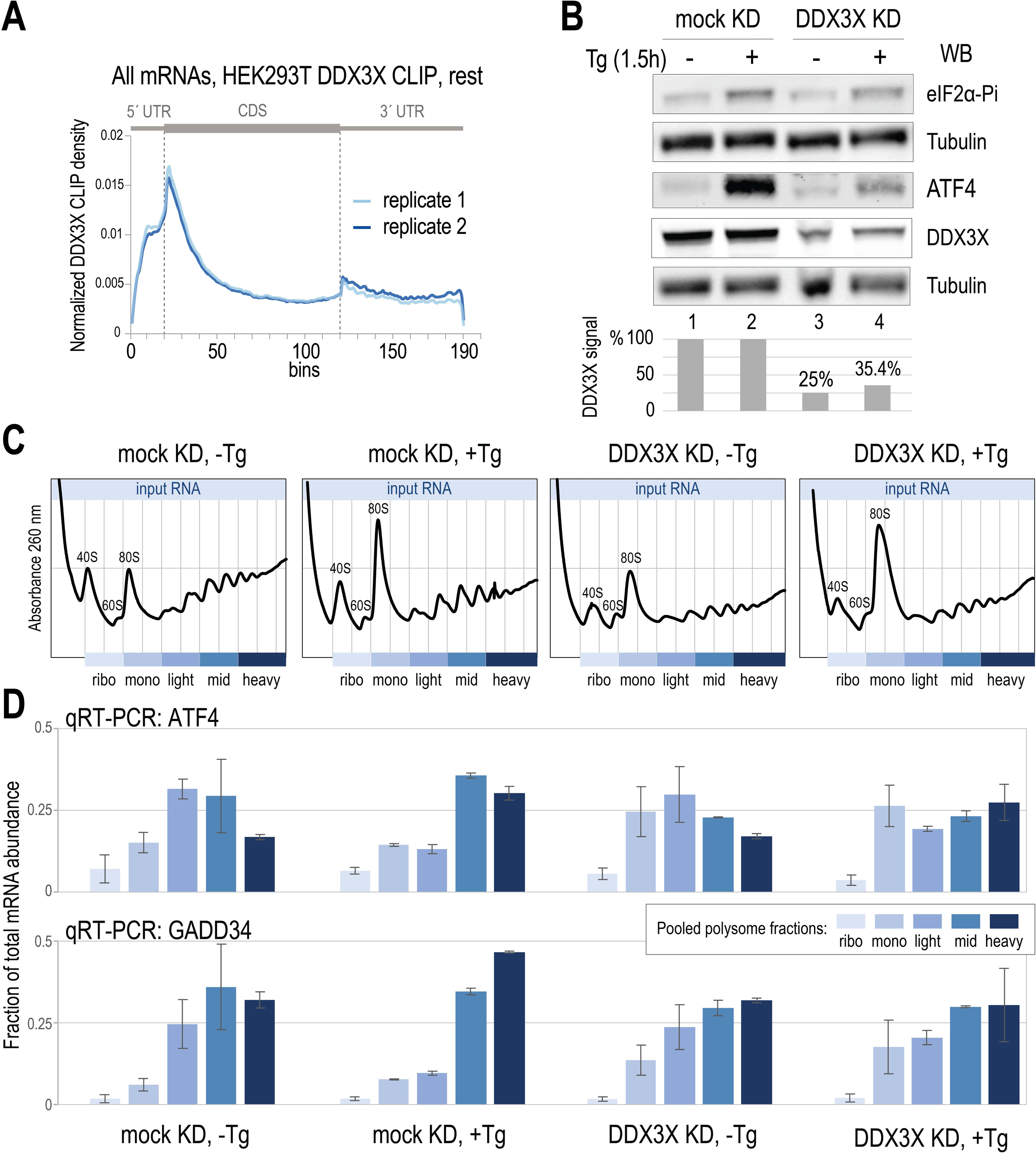
DDX3X CLIP-Seq in human cells, and the impact of Tg stress and DDX3X KD measured by polysome profiling. (A) DDX3X CLIP density across the meta mRNA in non-stressed 293T cells. (B) Western blot for indicated proteins in cell lysates used for polysome profiling, for indicated conditions. Lower panel: quantification of DDX3X signal from the same blot (vs tubulin). (C) 260 nm absorbance of polysome gradient collection, for indicated conditions. Vertical lines indicate individual fractions collected; blue colors indicate grouping of fractions for RT-qPCR analysis. (D) RT-qPCR analysis for ATF4 and GADD34 in grouped fractions as in (C), for indicated conditions. N=3, error bars: SD.

To functionally define DDX3X-regulated mRNAs, we knocked down (KD) DDX3X by RNAi in control and Tg-treated cells and performed polysome profiling. We verified ∼70% DDX3X protein depletion (Figure 2B) and used a non-targeting siRNA as negative control (mock). Cells were treated with Tg for 1.5 h to induce acute ER stress^35^, which was confirmed by increased eIF2α phosphorylation (Ser52) and ATF4 protein levels (Figure 2B). Notably, ATF4 protein was reduced in DDX3X KD cells under stress, consistent with previously reported DDX3X-dependent translation of ATF4 mRNA ^32,33^. We performed polysome profiling on 10–50% sucrose gradients in two biological replicates per condition: mock KD −Tg, mock KD +Tg, DDX3X KD −Tg, and DDX3X KD +Tg. Absorbance at 260 nm revealed the expected polysome reduction and 80S monosome increase upon Tg treatment in control cells. DDX3X KD reduced polysomes in non-stressed cells and, under ER stress, prevented further polysome collapse despite an increase in the 80S peak (Figure 2C). We extracted RNA from indicated grouped fractions (Figure 2C) and confirmed by RT-qPCR that ATF4 and GADD34 mRNAs shifted to heavy polysomes in control KD +Tg^35^, but not in DDX3X KD +Tg cells (Figure 2D), implicating DDX3X in stress-induced translational activation.

We then performed RNA-Seq on total (input) and heavy polysome RNA (highlighted dark blue, Figure 2C) from two replicates per condition. Plotting translation changes against mRNA abundance in mock KD +Tg revealed primarily translational (not abundance) shifts, consistent with MEFs, including upregulation of GADD34 and a trend for ATF4 (Figure 3A, Table S2).

**Figure 3.**
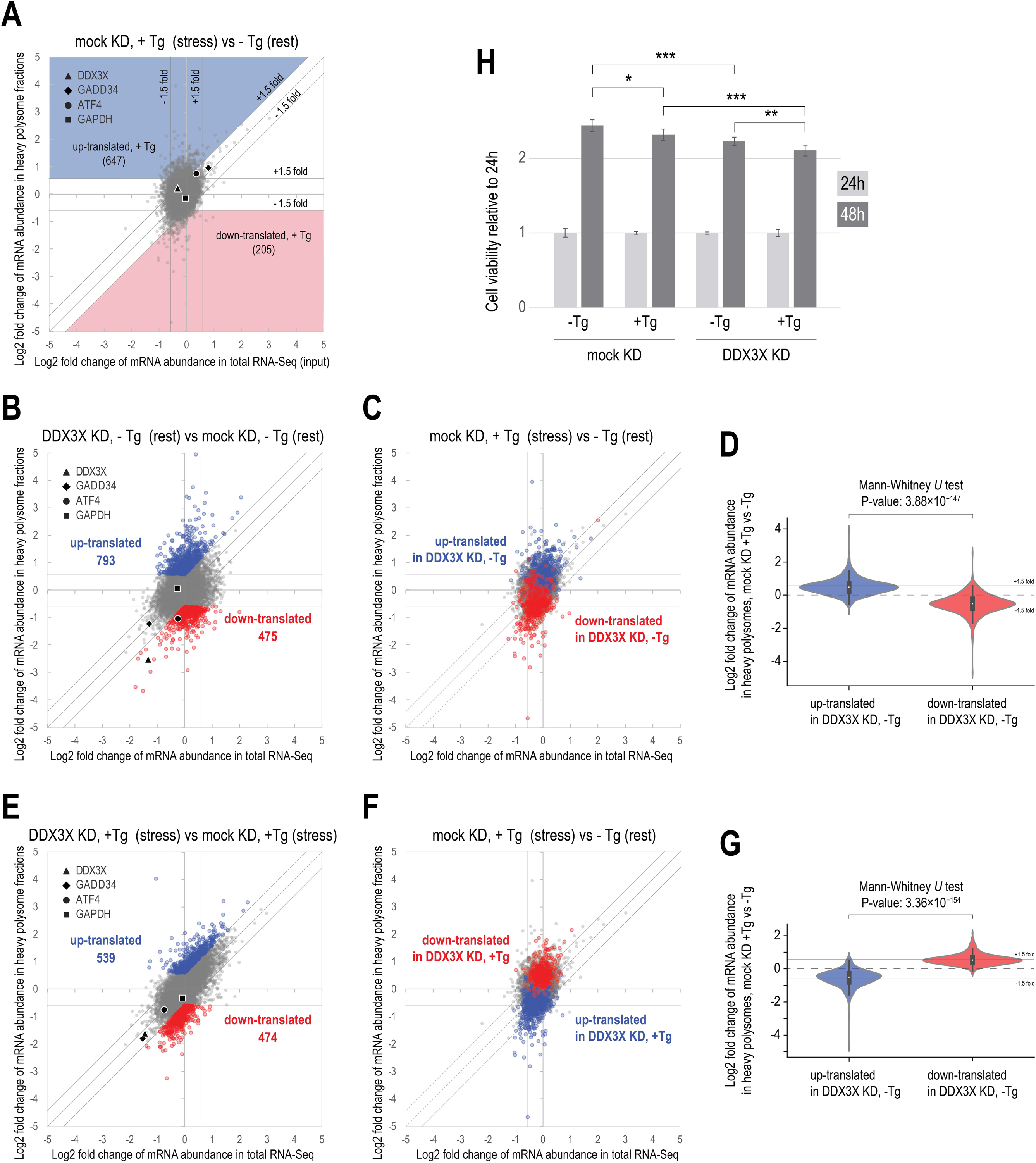
mRNA abundance and polysome association changes upon DDX3X KD in the absence or presence of Tg induced stress. (A) Polysome profiling mRNA scatterplot for changes in abundance and polysome association in mock KD 293FT cells treated with Tg compared to no Tg treatment. Four individual mRNAs are highlighted. (B) Polysome profiling mRNA scatterplot for changes in abundance and polysome association for DDX3X KD compared to mock KD, no Tg treatment. mRNAs showing 21.5 fold increased (up-translated) and decreased (down-translated) polysome association are highlighted blue and red respectively. (C) mRNA scatterplot as in (A) with mRNAs highlighted according to polysome association change as in (B). (D) Violin plot for highlighted mRNAs in (C) and statistic test as indicated. (E) Polysome profiling mRNA scatterplot for changes in abundance and polysome association for DDX3X KD compared to mock KD, after Tg treatment. Up-and down-translated mRNAs are highlighted blue and red respectively. (F) mRNA scatterplot as in (A) with mRNAs highlighted according to polysome association change as in (E). (G) Violin plot for highlighted mRNAs in (E) and statistic test as indicated. (H) Cell viability for mock and DDX3X KD cells following DMSO (control) or CPA treatment (1.5h and then wash-off), measured at 24h and 48h after CPA wash-off, relative to 24h. N=3, error bars: SD.

DDX3X KD in non-stressed cells resulted in increased heavy polysome association for 793 mRNAs (up-translated) and decreased association for 475 mRNAs (down-translated), including ATF4 (Figure 3B) and previously identified DDX3X-dependent mRNAs ODC1 and PRKRA (Table S2). To investigate how these dysregulated mRNAs behave in control cells under acute ER stress, we marked the DDX3X KD −Tg up-and down-translated mRNAs (Figure 3B) onto the polysome profiling scatterplot comparing mock KD +Tg vs. −Tg. We observed that the mRNAs up-translated in DDX3X KD under non-stressed conditions tend, on average, to also shift toward higher polysome association in mock KD +Tg cells (Figures 3C and 3D). Moreover, a statistically significant number of mRNAs (242) are up-translated more than 1.5x fold, both in DDX3X KD - Tg and in mock KD +Tg (hypergeometric test, p value: 1.12×10^−95^) (Figures 3D and S5A), indicating that DDX3X suppresses stress-responsive mRNAs in basal conditions. Over-represented gene ontology (GO) categories ^51^ include regulation of transcription, chromatin organization and remodeling, serine phosphorylation (among which WNK1, PRKCA, PRKCQ, PKN2), regulation of cAMP dependent protein Kinases (PRKAR1A, PRKAR2A, ATP2B4, RAPGEF2) and histone lysine demethylation (KDM6B, JMJD1C, KDM6A) (Figures S5A-S5D, Table S3).

Conversely, the 475 down-translated mRNAs in DDX3X KD −Tg were also on average down-regulated in mock KD +Tg, with 99 mRNAs showing ≥1.5-fold reduction in both (hypergeometric p = 1.83×10⁻⁷¹; Figures 3D, S5E). These represent transcripts positively regulated by DDX3X in non-stressed cells but suppressed during stress. GO terms included ribosome biogenesis (ribosomal proteins, known targets of DDX3X regulation ^42,52^), translation, ER targeting, and mitochondrial NADH dehydrogenase assembly (Figures S5E–S5H, Table S3).

We next examined DDX3X-regulated mRNAs in Tg-treated cells. DDX3X KD +Tg led to down-translation of mRNAs that are normally up-regulated in mock KD +Tg, and vice versa (Figures 3F, 3G), indicating that DDX3X shapes the stress-induced translational landscape. This was further supported by a significant number of mRNAs showing opposite translation shifts (≥1.5-fold change) in DDX3X KD +Tg and mock KD +Tg cells (Figures 3G, S6A, S6B). These included heat shock proteins and UGGT1, key factors in the response to misfolded proteins in the ER (’De Novo’ Post-Translational Protein Folding, GO:0051084), which were down-translated upon DDX3X KD but up-translated in control cells under stress.

GO terms enriched among mRNAs up-translated in mock KD +Tg and down-regulated in DDX3X KD +Tg included mRNA splicing, telomere maintenance (subunits of the chaperonin-containing T-complex, TRiC), RNA processing and stabilization, chaperone-dependent protein refolding, regulation of translation, RNA and DNA metabolic processes, and regulation of apoptosis, (Figures S6C-S6E, Table S3). Conversely, genes up-translated in DDX3X KD +Tg but suppressed in mock KD +Tg were enriched for processes such as mRNA translation (ribosomal proteins, which are down-translated in DDX3X KD-Tg), mitochondrial ATP synthesis coupled with electron transport (subunits of the NADH ubiquinone oxidoreductase mitochondrial complex I) and iron sulfur cluster assembly (ISCA2, BOLA3, CISD1), and SRP-dependent ER targeting (Figures S6F-S6H, Table S3).

To assess the physiological significance of the translational regulation by DDX3X in basal and stress conditions, we measured cell viability during recovery from acute ER stress by cyclopiazonic acid (CPA), a reversible ER stressor. Both CPA treatment and DDX3X KD individually reduced cell viability at 48h, and their combination led to the strongest effect (Figure 3H), supporting the view that DDX3X-mediated translational control promotes stress adaptation and survival, as suggested by our GO analysis.

Together, these findings support a model in which DDX3X acts as a dual-function regulator, enhancing or repressing translation depending on cellular context. In non-stressed cells DDX3X suppresses stress-activated mRNAs many of which are involved in transcriptional regulation and silencing, while promoting translation of core biosynthetic components such as ribosomal proteins. Under ER stress, this regulation is reversed: DDX3X promotes translation of chaperones and RNA processing factors while down-regulating energetically expensive processes, likely to aid adaptation and energy conservation.

### DDX3X CLIP-Seq abundance of an mRNA is not predictive of its regulation by DDX3X

We next sought to further explore our finding that DDX3X can both promote and suppress translation depending on mRNA context and stress conditions. We asked if DDX3X binding of mRNAs as represented by overall CLIP mRNA abundance (in cells-Tg) is correlated with the ratio of heavy polysomes over total RNA abundance (H/T ratio, as a measure of mRNA translation efficiency in a single condition), under the simple assumption that DDX3X binding drives regulation. We observed very weak positive correlation between DDX3X CLIP TPM (log10) and H/T ratio in mock KD-Tg cells for up-translated and down-translated mRNAs,-0.031 (R^2^: 0.0009) and-0.283 (R^2^: 0.0803) respectively (Figures S7A and S7B), which further supports the notion that a simple model relying on just the degree of DDX3X associating with an mRNA has very limited predictive value for mRNA translational regulation, if at all. Moreover, we see mRNAs with high and low translation efficiency being lowly or highly bound in both these categories; the same is true for all bound mRNAs (Figure S7C), and mRNAs that are bound but whose translation does not change when DDX3X is knocked down in-Tg cells (Figure S7D). These results support the notion that the degree to which DDX3X is associated with an mRNA across its entire length cannot predict if and how this mRNA is being regulated by DDX3X in human cells, in agreement with the analysis of DDX3X CLIP in MEFs. Nevertheless, DDX3X clearly selects a subgroup of the mRNAs that it binds, for positive and negative regulation of translation in non-stressed and stressed cells.

### mRNA binding and bidirectional translational regulation by DDX3X in the context of 48S scanning complex

Given that a substantial number of DDX3X-bound mRNAs are not dysregulated upon DDX3X knockdown (Figures 3B, 3E, and S7D), and that the extent of mRNA binding by DDX3X is not predictive of regulatory outcomes, it follows that the decision of whether and how an mRNA associated with DDX3X will be regulated, is made after RNA-protein association has occurred. We hypothesize that the initial engagement of DDX3X with its mRNA targets is not determined by DDX3X alone, but rather occurs in the context of the 48S scanning complex. Supporting this view, there is prior evidence of DDX3X associating with multiple translation initiation factors and related proteins^42,53–55^. We performed mass spectroscopy of peptide-eluted DDX3X IP samples from human cells, and identified the eIF3 complex (12 out of 13 subunits), a key component of the 43S PIC and the 48S scanning complex, as the top coimmunoprecipitated set of proteins with DDX3X (and absent from control IP), in an RNA-independent manner (Figure 4A, B; Table S4). Co-IP with eIF4G was also observed in the same experiment. We also independently verified the RNA-independent co-immunoprecipitation of DDX3X with eIF3D (Figure S8A). Moreover, we observed pronounced and enriched CLIP binding of DDX3X on helix 16 of the 18S rRNA, relative to RNA-Seq coverage of 18S and 28S rRNA (Figure S8B), consistent with previous observations ^13–15^. This provides evidence for a precise positioning of DDX3X within the 48S scanning complex, specifically at the mRNA entry site of the 40S ribosomal subunit. Notably, at the moment of start codon decoding by the 48S complex, this positioning places DDX3X immediately downstream of the start codon, within the early coding sequence (CDS) ^1^. These findings support a model in which DDX3X is recruited to the 48S complex through its interaction with eIF3, enabling it to participate in 5‘ UTR scanning and translation initiation.

**Figure 4.**
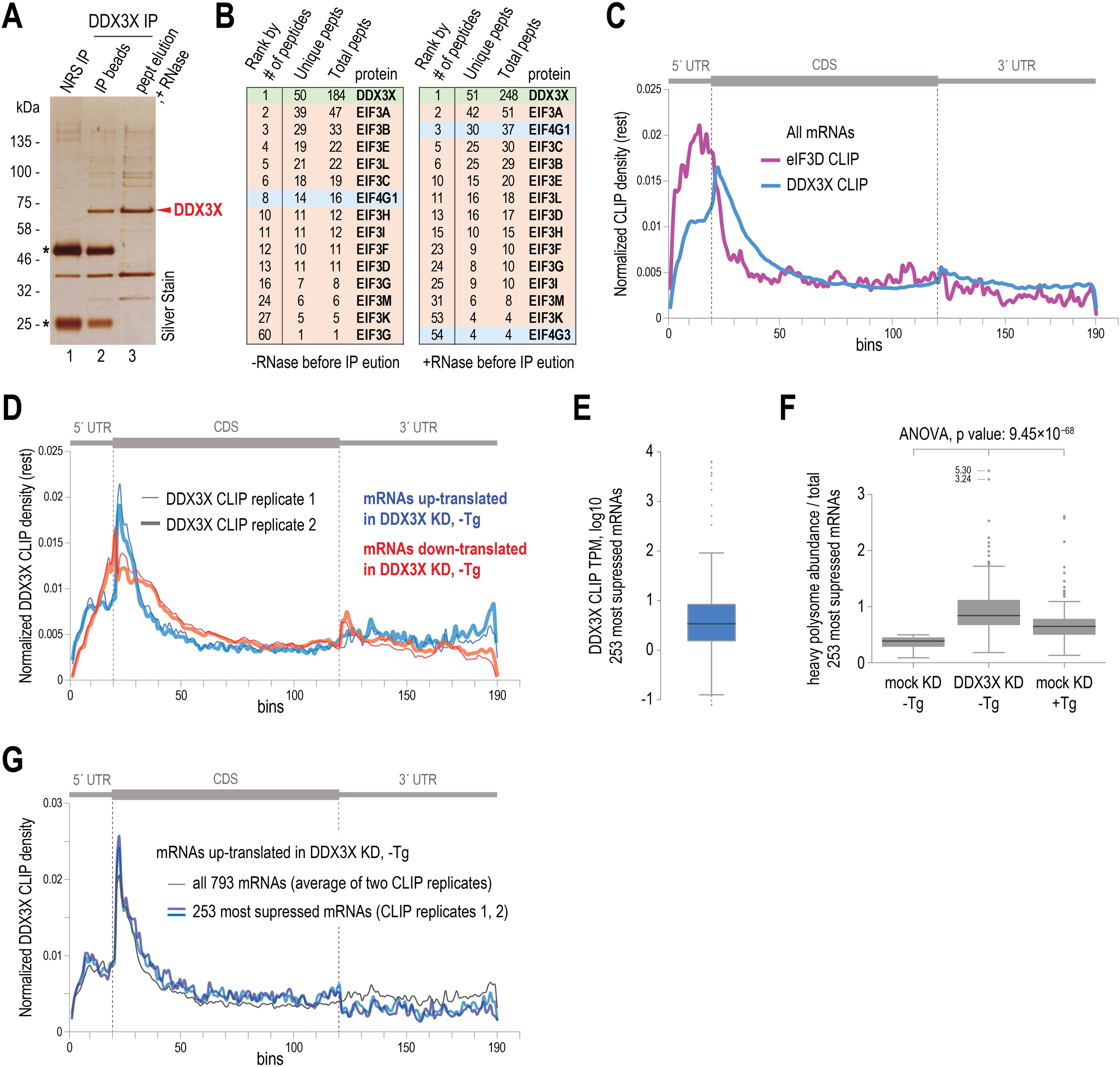
Association of DDX3X with eIF3 and DDX3X regulation of mRNAs in the context of 48S scanning process. (A) Silver stain of DDX3X IP from human cells. Major immunoprecipitated proteins are marked. Lane 1: negative control IP with normal rabbit serum (NRS IP). Lane 2: reducing elution of DDX3X IP beads. Lane 3: peptide elution of DDX3X IP, after treatment with RNase A. *: heavy and light Ab chain. (B) Number of unique and total peptides (pepts) of eIF3 subunits and eIF4G proteins, identified by mass spec of gel section containing the entire peptide-eluted DDX3X IP sample. The entire list of peptides can be found in Table S4. (C) DDX3X CLIP density on all mRNAs for eIF3D (magenta) and DDX3X CLIP (blue) in-Tg cells. (D) DDX3X CLIP density on up-(blue tones) and down-translated mRNAs (red tones) in DDX3X KD-Tg. (E) DDX3X CLIP abundance (log10 TPM) of 253 mRNAs with H/T ratio <0.5 in-Tg mock KD cells (see also Figure S7A) that are also up-translated in DDX3X KD-Tg. (F) Box plots of H/T ratios for the same 253 mRNAs in indicated conditions. Two-tailed t-test comparisons between-Tg mock and DDX3X KD (p-value: 6.1×10^-22^); mock KD-Tg and mock KD +Tg (p-value: 1.1×10^-19^), and three-way ANOVA test as indicated. (G) DDX3X CLIP density (-Tg) on all 793 up-translated mRNAs in DDX3X KD-Tg (thin line) and 253 most suppressed mRNAs (blue tones).

To further support this view, we performed eIF3D CLIP (Figure S8C) and compared the eIF3D and DDX3X CLIP densities across the meta-mRNA (Figure 4C). eIF3D CLIP density is enriched throughout the 5‘ UTR peaking upstream of the start codon, and dropping sharply thereafter, consistent with rapid eIF3 release following initiation^56^. In contrast, DDX3X CLIP density gradually increases across the 5‘ UTR, peaks just after the start codon, and persists into the CDS, remaining detectable beyond the eIF3D signal (Figure 4C). This pattern suggests a role for DDX3X in early translation elongation in addition to 5‘UTR scanning. Supporting this, we detected DDX3X in heavy polysome fractions (Figure S8D). Together, these findings suggest that during 5‘ UTR scanning by the 48S complex, eIF3D reaches maximal recruitment earlier than DDX3X, consistent with our model that DDX3X depends on eIF3 association for 48S recruitment. Furthermore, the distinct CLIP coverage patterns reinforce our conclusion that DDX3X’s presence in the early CDS is critical for mRNA translation regulation.

We reasoned that if DDX3X engages mRNAs as part of the 48S scanning complex, and if regulatory decisions for promoting or suppressing translation are made during this process, this should be reflected in the meta-mRNA DDX3X CLIP density. Indeed, mRNAs that are up-translated or down-translated upon DDX3X KD (-Tg), as well as those showing no change in polysome association, all exhibit enrichment of DDX3X binding around the start codon (Figure 4D and Figure S9A), consistent with our model. Notwithstanding the overall similarity, we observed distinct binding profiles of DDX3X on transcripts whose translation is either stimulated or suppressed by this helicase (Figure 4D). mRNAs down-translated upon DDX3X KD, indicating positive regulation by DDX3X, exhibited gradually increasing binding in their 5‘ untranslated regions (5‘ UTRs) (Figure 4D, red trace). In contrast, mRNAs up-translated upon DDX3X KD, suggesting suppression by DDX3X, showed an abrupt increase of density downstream of the start codon in the early CDS (Figure 4D, blue trace). These distinct binding patterns suggest that DDX3X engages in at least two mechanistically distinct modes of translational regulation.

While it is expected that positive regulation of mRNA translation by DDX3X takes place during 48S scanning and translation initiation, our analysis suggests that DDX3X-dependent mRNA suppression is also enacted in the same context. To further explore this, we looked into the up-translated mRNAs in DDX3X KD-Tg and focused on the 253 mRNAs with the lowest H/T ratio (<0.5) in mock KD-Tg (Figure S7A). These mRNAs are well bound by DDX3X (median log10 CLIP TPM: 0.532) (Figure 4E), depleted from polysomes in control cells (median H/T ratio: 0.387), and are robustly up-translated in DDX3X KD cells (median H/T: 0.840, a ∼2.2x increase, Figure 4F), suggesting that they are actively suppressed by DDX3X in non-stressed cells. Notably, these 253 mRNAs are also significantly up-translated in mock KD +Tg (median H/T: 0.648; Figure 4F), implying that this suppression mechanism is pre-emptively engaged under basal conditions and lifted in response to ER stress. The DDX3X CLIP density for this subset also shows strong enrichment directly downstream of the start codon, closely matching that for all 793 up-translated mRNAs (Figure 4G).

These findings support a model in which DDX3X contacts mRNAs predominantly through its incorporation into the scanning 48S complex rather than through intrinsic recognition of target transcripts. Some of the mRNAs undergoing scanning rely on DDX3X to promote or repress their translation, while others do not, and for these, DDX3X binding has no functional consequence.

### Distinct mechanisms of mRNA regulation by DDX3X revealed by initiation profiling

The prevailing model of DDX3X translation regulation posits that DDX3X promotes translation by resolving secondary structures in the 5‘UTR. To test this model and investigate the biological context of DDX3X-dependent regulation under basal and ER stress conditions, we focused on a core subset of mRNAs exhibiting a bidirectional switch in translation, as a result of DDX3X KD and the presence of ER stress. Specifically, we identified two groups: (1) mRNAs down-translated in DDX3X KD under non-stress conditions and up-translated in DDX3X KD during acute ER stress; and (2) mRNAs which show the opposite behavior (Figure 5A). As previously noted (Figure 3), translation of many of these transcripts is also regulated in ER stress in mock KD cells: those down-translated upon DDX3X KD in non-stressed cells are also down-translated in mock KD during ER stress, and those up-translated upon DDX3X KD under basal conditions are also up-translated in mock KD following stress (Figures S5, S6). Therefore, we delimited the two core groups as follows: 63 mRNAs designated as “DDU” (<u>D</u>own in DDX3X KD-Tg ∩ <u>D</u>own in mock KD +Tg ∩ <u>U</u>p in DDX3X KD +Tg) and 32 mRNAs designated as “UUD” (<u>U</u>p in DDX3X KD-Tg ∩ <u>Up</u> in mock KD +Tg ∩ <u>D</u>own in DDX3X KD +Tg) (Figure 5A). STRING analysis^57^ of the “DDU” mRNAs revealed a significant enrichment of physical and functional interactions between them (PPI enrichment p < 1×10⁻¹⁶), with prominent overrepresentation of translation-related processes, including ribosomal protein synthesis, co-translational targeting to ER, and formation of free 40S ribosome subunits (Figure 5B). Additional enrichment in mitochondrial functions and respiratory electron transport suggests potential coordinated regulation of translational control and metabolic adaptation by DDX3X. “UUD” mRNAs are significantly enriched for RNA and nucleic acid binding functions, forming a network of mRNA processing factors (Figure 5C). These two groups represent core subpopulations of DDX3X-regulated mRNAs that are bidirectionally responsive to DDX3X regulation and play roles in the cellular response to stress.

**Figure 5.**
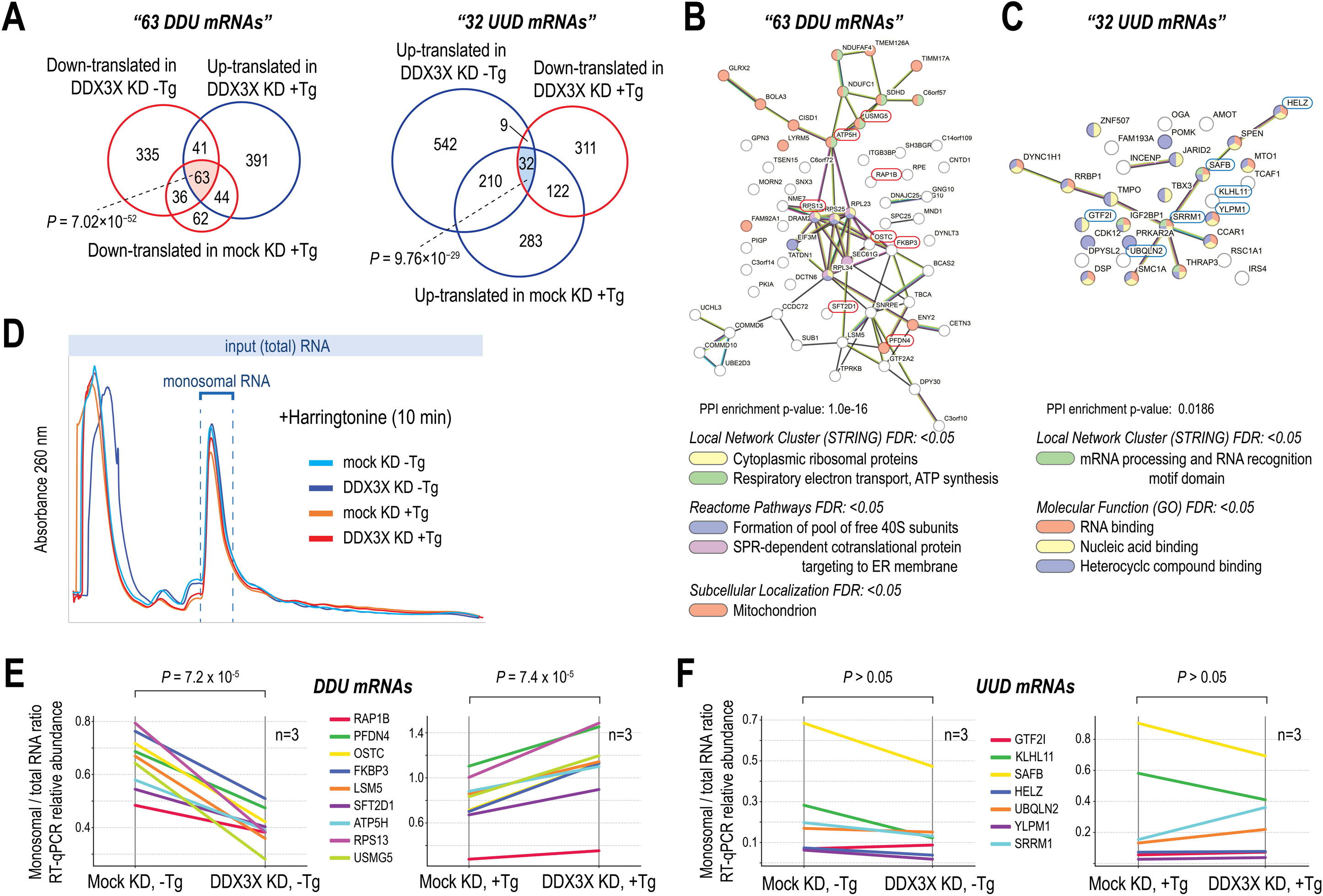
Different mechanisms of translational regulation of two core subsets of DDX3X target mRNAs (A) Venn diagrams of DDX3X regulated mRNAs and statistic test (hypergeometric test) for the overlaps as indicated. The triple overlap is highlighted blue for 63 “DDU” mRNAs and red for 32 “UUD” mRNAs respectively. (B) STRING database analysis for “DDU” mRNAs. Statistically enriched terms (FDR< 0.05) are shown; mRNAs are colored by term enrichment. mRNAs selected for further analysis below are outlined in red. (C) STRING database analysis for “UUD” mRNAs. Statistically enriched terms (FDR< 0.05) are shown; mRNAs are colored by term enrichment. mRNAs selected for further analysis below are outlined in blue. (D) 260 nm absorbance of polysome gradients after harringtonine treatment for indicated conditions. RNA was extracted from the monosomal peak and from total input RNA. (E) Monosome-to-total (input) mRNA abundance ratios were measured for “DDU” mRNAs by RT-qPCR following polysome profiling with harringtonine treatment, which captures newly initiated ribosomes. Each line connects the mean values of the ratio for an individual mRNA in mock KD and DDX3X KD conditions. Significance was evaluated with paired t-test, n=3. (F) Same as in (E) for “UUD” mRNAs.

To determine whether DDX3X regulates these mRNAs at the level of translation initiation, we performed polysome profiling following harringtonine (HRN) treatment (Figure 5D). HRN binds to the ribosomal A-site and prevents formation of the first peptide bond, effectively arresting translation immediately after initiation. Importantly, HRN does not interfere with 48S scanning, start codon decoding, or elongation by pre-engaged ribosomes. We collected HRN-treated polysome profiles in biological triplicates, isolated total (input) RNA and monosomal RNA fractions (Figure 5D), and performed RT-qPCR (equal mass of RNA was used across samples for RT) to determine the ratio of monosomal RNA to total RNA for each target transcript, which represents the accumulation of each mRNA in the monosomal fraction during HRN treatment, effectively representing the rate by which the mRNA is initiated in each condition (Figure 5E). We observed that “DDU” mRNAs show significantly reduced initiation in non-stressed cells upon DDX3X KD compared to mock KD (paired t-test, P = 7.2 × 10⁻⁵), and significantly increased initiation during ER stress upon DDX3X KD (paired t-test, P = 7.4 × 10⁻⁵), thereby supporting a role for DDX3X in regulating translation initiation through its binding in the 5‘ UTR of “DDU” mRNAs (Figure 5E), which are involved in energy metabolism and the translation machinery. Conversely, “UUD” mRNAs did not show a statistically significant change in monosomal/total RNA ratios in DDX3X KD compared to mock KD, or a concerted change thereof in stress (Figure 5F), suggesting that DDX3X regulates “UUD” mRNAs through a distinct, likely a mechanism that involves regulation after initiation. Taken together with the observed differences in DDX3X distribution around the start codon for up-and down-translated mRNAs, the above suggest that DDX3X employs mechanistically distinct regulation for these two mRNA groups.

### N4-acetylation of cytidines is an mRNA feature associated with DDX3X-mediated translational suppression

We hypothesized that during 48S scanning, DDX3X interacts with local mRNA features that influence whether it engages productively with its targets. In the absence of such features, transient association may be functionally neutral, which is consistent with the observation that a large group of DDX3X-bound mRNAs are not regulated upon knockdown. Given that DDX3X binding peaks around the start codon and shows prominent differences for groups of mRNAs that are differentially regulated by DDX3X (Figure 4D), we reasoned that critical regulatory features may reside in this region and contribute to differential outcomes.

Recent studies have implicated N4-acetylcytidine (ac4C), catalyzed by NAT10, as a post-transcriptional mark that modulates translation initiation in a position-dependent manner ^46,47,58^. We hypothesized that ac4C may serve as such a feature, influencing DDX3X-mediated regulation. To test this, we examined published ac4C FAM-Seq and RNA-IP-Seq data from 293T cells ^59^, and observed distinct patterns in the distribution of ac4C-modified reads near the start codon of DDX3X target mRNAs (Figure 6A). Transcripts down-translated upon DDX3X KD (i.e., normally promoted by DDX3X), for which we observed control at the level of initiation exhibited increasing ac4C density in the 5′ UTR with a peak at the start codon. In contrast, mRNAs up-translated in DDX3X KD (normally repressed by DDX3X) were depleted of ac4C in the 5′ UTR but showed elevated ac4C density extending into the CDS.

**Figure 6.**
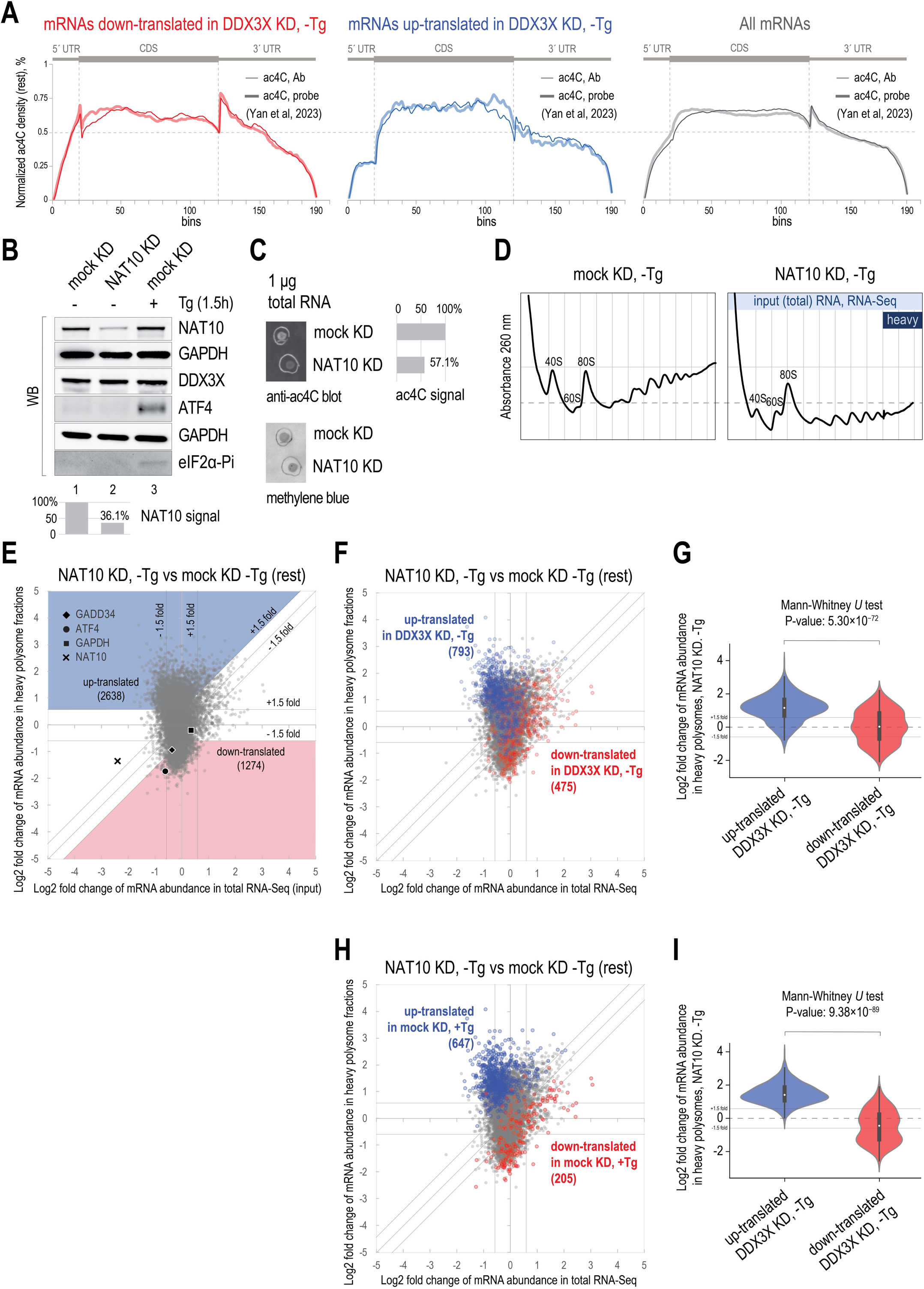
mRNA co-dependence on DDX3X and ac4C modification for translational regulation. (A) ac4C density across the meta mRNA in DDX3X KD-Tg cells, for down-translated mRNAs (left, red), up-translated mRNAs (middle, blue) and all mRNAs (right, grey). Data from Yan et al., 2023, showing two separate ac4C-Seq methods that the authors employed, one using an Ab against ac4C, and a second one using a chemical probe. (B) Western blot for indicated proteins in in mock and NAT10 KD cell lysates, as indicated. Lower panel: quantification of NAT10 signal from the same blot (vs GAPDH). (C) Immuno-dot blot for ac4C using mock and NAT10 KD total RNA, top panel. Methylene blue stain, bottom panel, was used as a loading reference. Ac4C signal quantification is shown to the right of the top panel, one representative experiment shown. (D) 260 nm absorbance of polysome gradient collection, for indicated conditions. Vertical lines indicate individual fractions collected; blue colors indicate grouping of fractions for RNA-Seq library preparation. (E) Polysome profiling mRNA scatterplot for changes in abundance and polysome association in NAT10 KD compared to mock KD cells,-Tg. Four individual mRNAs are highlighted. (F) Polysome profiling mRNA scatterplot as in (E) with mRNAs highlighted according to polysome association change as in DDX3X KD-Tg (Figure 3B). (G) Violin plot for highlighted mRNAs in (F) and statistic test as indicated. (H) Polysome profiling mRNA scatterplot as in (E) with mRNAs highlighted according to polysome association changes in mock KD +Tg vs-Tg (Figure 3A). (I) Violin plot for highlighted mRNAs in (H) and statistic test as indicated.

To explore this relationship experimentally, we depleted NAT10 by RNAi in 293FT cells (−Tg) and performed polysome profiling (two replicates). NAT10 KD reduced protein levels by ∼65%, while DDX3X levels remained unchanged (Figure 6B); we also verified reduced total RNA acetylation by immuno-dot blot (Figure 6C). Polysome absorbance profiles showed a pronounced reduction in polysomes (Figure 6D), and mRNA-level polysome profiling revealed widespread changes in translational efficiency (Figure 6E), consistent with a global role for ac4C in translation regulation. Overlaying DDX3X-regulated mRNAs onto the NAT10 KD data showed that transcripts up-translated upon DDX3X KD were also strongly enriched among up-translated mRNAs in NAT10 KD (598 of 793 with ≥1.5-fold increase; Figures 6F, 6G). This overlap suggests that efficient translational repression of these targets depends on both DDX3X and ac4C in non-stressed cells. In contrast, down-translated mRNAs in DDX3X KD showed variable responses to NAT10 KD, indicating additional regulatory layers. Interestingly, up-and down-translated mRNAs during ER stress in mock KD cells also appear regulated in the same directions in NAT10 KD-Tg (Figures 6H and 6I), suggesting that ac4C contributes more broadly to the regulation of stress-responsive mRNAs, for which DDX3X is also responsible. However, the greater number of affected transcripts in NAT10 KD relative to DDX3X KD implies that ac4C is involved in additional regulatory mechanisms beyond those governed by DDX3X.

These results suggest that ac4C modification patterns near the start codon contribute to the selective engagement of DDX3X with its targets, particularly in translational repression under non-stressed conditions, and may play a broader role in coordinating mRNA translation during stress response.

## Discussion

Our study provides compelling evidence that DDX3X serves as a regulatory switch in mRNA translation, orchestrating a dynamic response to acute cellular stress. We show that DDX3X, as a key component of the eIF4F/48S scanning complex, can act as both a translational activator and repressor in both non-stressed and stressed cells, with its regulatory functions consistent with unique mRNA features, including cytidine acetylation patterns around the start codon. This positions DDX3X as a versatile mediator of cellular response to stress.

Focusing on mRNAs showing only heavy polysome association changes in our polysome profiling experiments, we observed that DDX3X appears to act as a stress sensor that modulates translation, likely to adapt cellular energy demands and ribosome biogenesis. During acute ER stress, DDX3X shifts to promote the translation of chaperones and stress response factors while suppressing translation-related mRNAs, such as ribosomal proteins, whose translation DDX3X promotes in non-stressed cells. In non-stressed cells, DDX3X represses stress-activated mRNAs, including those involved in transcriptional regulation and chromatin remodeling. This dual regulatory activity suggests that DDX3X may be effectively “priming” the cell for a rapid reaction and adaptation to acute episodes of ER stress. We find that DDX3X KD reduces cell survival especially in ER stress, supporting the notion that DDX3X regulates mRNAs to enhance cellular fitness prior and in response to cellular stress. Our findings also align with observations in yeast, where the Hinnebusch group demonstrated a significant overlap between mRNAs down-regulated in a Ded1p mutant in non-stressed conditions, and mRNAs downregulated in WT cells exposed to glucose stress, suggesting a potential conserved function of this family of RNA helicases across species ^60^.

Despite select mRNA translational control, in non-stressed and stressed cells, CLIP-Seq revealed association of DDX3X with the vast majority of the mRNAs expressed in human and mouse cell lines, including mRNAs whose translation is not appreciably changed in acute ER stress, or when DDX3X is knocked down in non-stressed and stressed cells. Moreover, our data suggest that DDX3X does not dissociate from its mRNA targets transcriptome-wide in stress.

Our findings show a strong, RNA-independent, association of DDX3X with the eIF3 complex, consistent with previous observations^53,54^. DDX3X and eIF3d CLIP-Seq meta mRNA densities suggest DDX3X reaches mRNA recruitment maximum downstream of eIF3. Moreover, DDX3X shows enriched binding to helix 16 of the 18S rRNA at the mRNA entry site of the ribosome, which aligns with its proposed positioning for efficient scanning and start codon decoding. Taken together these observations lead us to hypothesize that DDX3X mRNA binding occurs in the context of eIF4F/48S scanning and not by intrinsic recognition, in agreement with a recent report that the interaction of DDX3X with the ribosome is necessary for mRNA regulation^43^. It is therefore likely that mRNA features encountered by DDX3X during scanning, dictate DDX3X action and, depending on the feature and its location result in translational stimulation or suppression (Figure 7).

**Figure 7.**
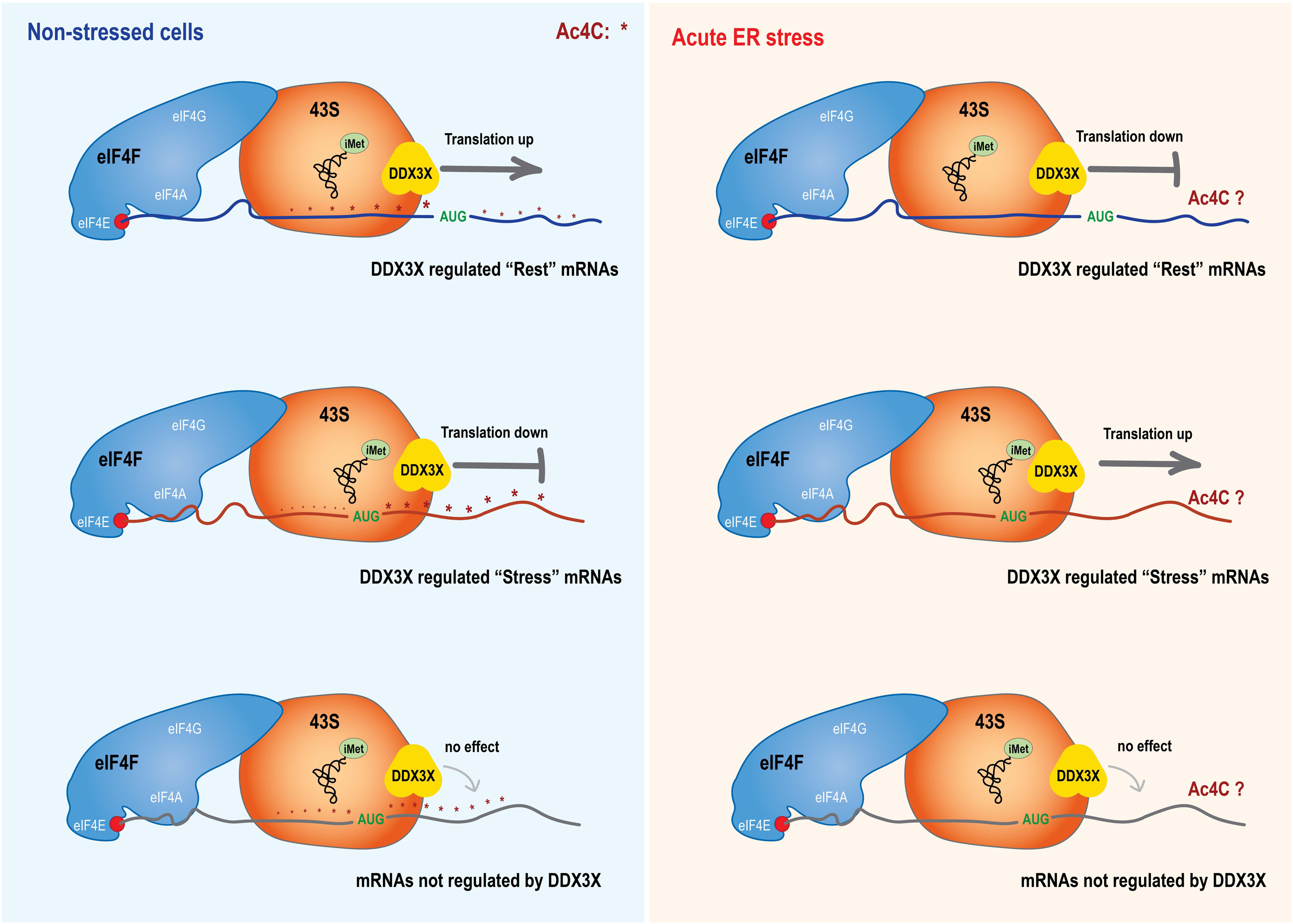
A model of DDX3X function as a dual switch regulator in translation initiation in non-stressed cells and cells during acute ER stress. “Rest” mRNAs denote mRNAs that are translated in non-stressed cells which show a DDX3X-dependent switch in translation in cells during acute ER stress; “Stress” mRNAs show opposite DDX3X-dependent regulation in the same conditions. The positioning of DDX3X relative to the start codon represents the peak CLIP density for the mRNAs in these categories.

The DDX3X CLIP-Seq density maps show a pronounced enrichment around the start codon, consistent across mouse and human cells, also observed in previous CLIP-Seq studies^14–16,42^. Notably, when CLIP meta mRNA plots are split into groups based on their regulation upon stress (Figures 1F and G), evidence on their mode of regulation is revealed by density pattern differences, previously masked in the all-mRNA meta plot. Stress related translation suppression results in reduced DDX3X presence in the 5‘UTR and especially downstream of the start codon followed by increased presence in the CDS, which would be consistent with futile mRNA scanning in stress (Figure 1G). On the other hand, up-translated mRNAs in stress show little DDX3X density change, suggesting stimulation by more efficient competition for the limited amounts of TC complex, as previously proposed ^30^.

Similarly, mRNAs differentially regulated by DDX3X in non-stressed cells show distinct DDX3X CLIP density, primarily around the start codon. mRNAs that depend on DDX3X for translation (down-translated upon DDX3X KD) show gradually increasing 5‘ UTR presence that peaks near the start codon, while mRNAs up-translated in DDX3X KD show a strong peak downstream of the start codon (Figure 4D). Importantly, by evaluating initiation rates with the use of harringtonine polysome profiling to measure the monosomal/total RNA enrichment, we find that mRNAs in the first category that also show opposite regulation in DDX3X KD in stressed cells (“DDU” mRNAs) are regulated at the level of translation initiation, while “UUD” mRNAs are not, suggesting distinct mechanisms of regulation by DDX3X for these two groups of transcripts. These observations lead us to hypothesize that key mRNA features involved in bidirectional regulation by DDX3X are located in the 5‘ UTR and the early CDS, focused around the start codon.

We focused on a recently identified mRNA PTM, ac4C, involved in mRNA translation regulation in a manner that depends on its location relative to the start codon ^47^. Importantly, ac4C density ^59^ shows distinct enrichment patterns for mRNAs differentially regulated by DDX3X (Figure 6A). Specifically, mRNAs that are translationally stimulated by DDX3X show increasing binding in their 5‘ UTR, and a similar pattern of ac4C density, while suppressed mRNAs show increased DDX3X binding downstream of the start codon and 5‘UTR ac4C depletion. Importantly, by profiling translation initiation using harringtonine, we find that mRNAs in the former category are regulated at the level of initiation, while mRNAs in the latter category are not. These results hint at a causative link between ac4C position on the mRNA and DDX3X regulation. We showed that mRNA suppressed by DDX3X are up-translated in NAT10 KD, demonstrating a possible co-dependence on both DDX3X and ac4C for this type of regulation in non-stressed cells, which suggests DDX3X might be a reader of post-transcriptional modifications. This interplay between ac4C modifications and DDX3X activity adds a new layer of complexity to our understanding of translational regulation. Although the 293T ac4C data we used in our analysis ^59^ are not nucleotide-resolved, distinct ac4C patterns around mRNA start codons are observed. Further work is needed to identify^61^ and induce^62^ ac4C modifications at precise locations and to determine if these can affect mRNA secondary structure formation, which in turn could be the substrate for DDX3X regulation ^45^. Recent findings by Fujita and colleagues have revealed that additional mRNA base modifications contribute to the regulation of translation initiation^63^. Specifically, 5-methylcytosine (5mC) and pseudouridine (Ψ) at near-cognate non-AUG start codons modulate their potency as translation initiation sites by tuning the base-pairing dynamics with the initiator tRNA. These findings further emphasize the complexity of epitranscriptomic regulation and support the role of diverse base modifications as dynamic elements of control of gene expression at the level of mRNA translation.

While translational stimulation by DDX3X is well documented, DDX3X translational suppression is more contested^17^. We provide evidence that mRNA features downstream of the start codon may be involved, in the comparison of the DDX3X CLIP meta plot for up-and down-translated mRNAs upon DDX3X KD. We observe a prominent peak of DDX3X density in the first 10 bins of the CDS in the former (Figure 4D), and which becomes even more prominent for the 253 most suppressed mRNAs by DDX3X (Figure 4G). One possible explanation is that DDX3X acts as a clamp on the mRNA preventing translation elongation. Long lived protein-RNA complexes, a.k.a. clamping, of Ded1 on mRNAs has been proposed and shown *in vitro* using non-hydrolysable ATP analogs ^64^, and a mechanism for mRNA clamping by eIF4AIII, a DEAD-box RNA helicase subunit of the Exon Junction complex, has been shown in structural studies ^65^; it remains to be seen if DDX3X can act as a clamp to suppress mRNA translation. It is also worth noting that, considering the proposed positioning of DDX3X downstream of the start codon during initiation, the binding pattern extending into the early CDS aligns with the role of DDX3X in 80S ribosome assembly ^8,54^ and potentially in early translation elongation, as suggested for Leishmania DDX3X ^66^.

The bidirectional regulation of mRNA translation by DDX3X has significant implications for understanding its role in cancer. Aberrant translation regulation, such as the dysregulation of mitochondrial enzymes and components of the respiratory chain, has been linked to oncogenic processes ^67^. Moreover, a role for DDX3X in aberrant ER stress in cancer cell lines has been shown ^41,42^. Our findings suggest that DDX3X-mediated translational control could be a factor in cancer cell survival and adaptation to acute episodes of stress ^68^, offering new avenues for potential therapeutic approaches.

Overall, our findings illuminate DDX3X’s central role as a mediator of translational regulation in stress, integrating RNA helicase function and post-transcriptional modifications. This dual regulatory capability positions DDX3X as a potential target for therapies aimed at diseases characterized by impaired stress responses, such as cancer and neurodevelopmental disorders.

## Limitations of the study

While our work advances the understanding of CLIP-Seq meta gene analysis by splitting the mRNAs in functionally coherent groups of mRNAs based on the results of KD experiments, it is still very challenging to derive precise molecular roles of an RBP by CLIP density meta-analysis, especially for proteins that participate in a dynamic process such as 48S scanning and initiation. In the case of DDX3X, the patterns of meta-mRNA CLIP-Seq densities depend on 48S scanning kinetics and 80S assembly, as well mRNA features such as secondary structures resolved by the helicase activity. As the atlas of mRNA features (sequence, structure, PTMs) is continuously populated, combined molecular, biochemical and machine learning approaches will be needed to build predictive models that can interpret CLIP-Seq data and derive biochemical and molecular activity from meta-mRNA CLIP-Seq density plots. To monitor translation efficiency in this study, we deep sequenced and compared input (total cytoplasmic mRNA), and mRNA associated with heavy polysomes; a more accurate picture could be drawn by sequencing additional gradient fractions including light and medium heavy polysomes, which however would greatly increase the complexity of the analysis. Additionally, our threshold for defining regulated mRNAs (1.5-fold change) may have excluded subtle yet biologically relevant changes. Finally, our NAT10 KD approach cannot distinguish the possible effects of ac4C reduction on rRNA (positions 1337 and 1842 of 18S are ac4C modified), or mRNA; however rRNA has significantly longer half life than mRNA, and within the time frame of the KD (48h) we expect that reduction of mRNA ac4C has the predominant effect.

## Data and code availability

RNA-seq and CLIP-Seq data have been deposited at GEO with accession numbers GSE282240, GSE282243 and GSE282245.Bioinformatic scripts created and used in this study are available at: https://github.com/Abdelmonsif/DDX3X

## Acknowledgments

We thank members of Vourekas and Hatzoglou laboratories for discussions and technical assistance. We thank Dr Jan Oppelt (University of Pennsylvania) and Dr Panagiotis Alexiou (Malta University) for initial bioinformatic analysis of an earlier version of this manuscript. We are grateful to University of Pennsylvania Center for AIDS Research (CFAR) Director Dr Ronald Collman, CFAR member Dr Una O’Doherty (now at Emory University), and Dr Zissimos Mourelatos (University of Pennsylvania) for their support at an initial stage of this project. This work was supported by Penn CFAR Pilot Grant AI045008, LSU Provost’s Fund for Innovation in Research, Louisiana Board of Regents RCS LEQSF (2023-26)-RD-A-16, and LSU start-up funds to A.V.; LSU Agricultural Center Collaborative Research Program (PG010315) to C.A.S. and A.V.; DK060596 to M.H.

## Author Contributions

A.V. conceived and supervised the project. A.V., M.H and C.A.S. collected financial support.

A.V. and A-E.M.S. designed experiments, and A-E.M.S. executed polysome profiling experiments with help from A.S. A-E.M.S. performed the bioinformatic analysis. J.M., K.A. assisted with experiments, M.D. assisted with bioinformatic analysis. M.H. contributed to experimental design and data interpretation. A.V. wrote the manuscript, and A-E.M.S, C.A.S. and M.H edited the manuscript.

## Declaration of interests

The authors declare no competing interests.

## Supplemental Figures and Legends

**Figure S1.**
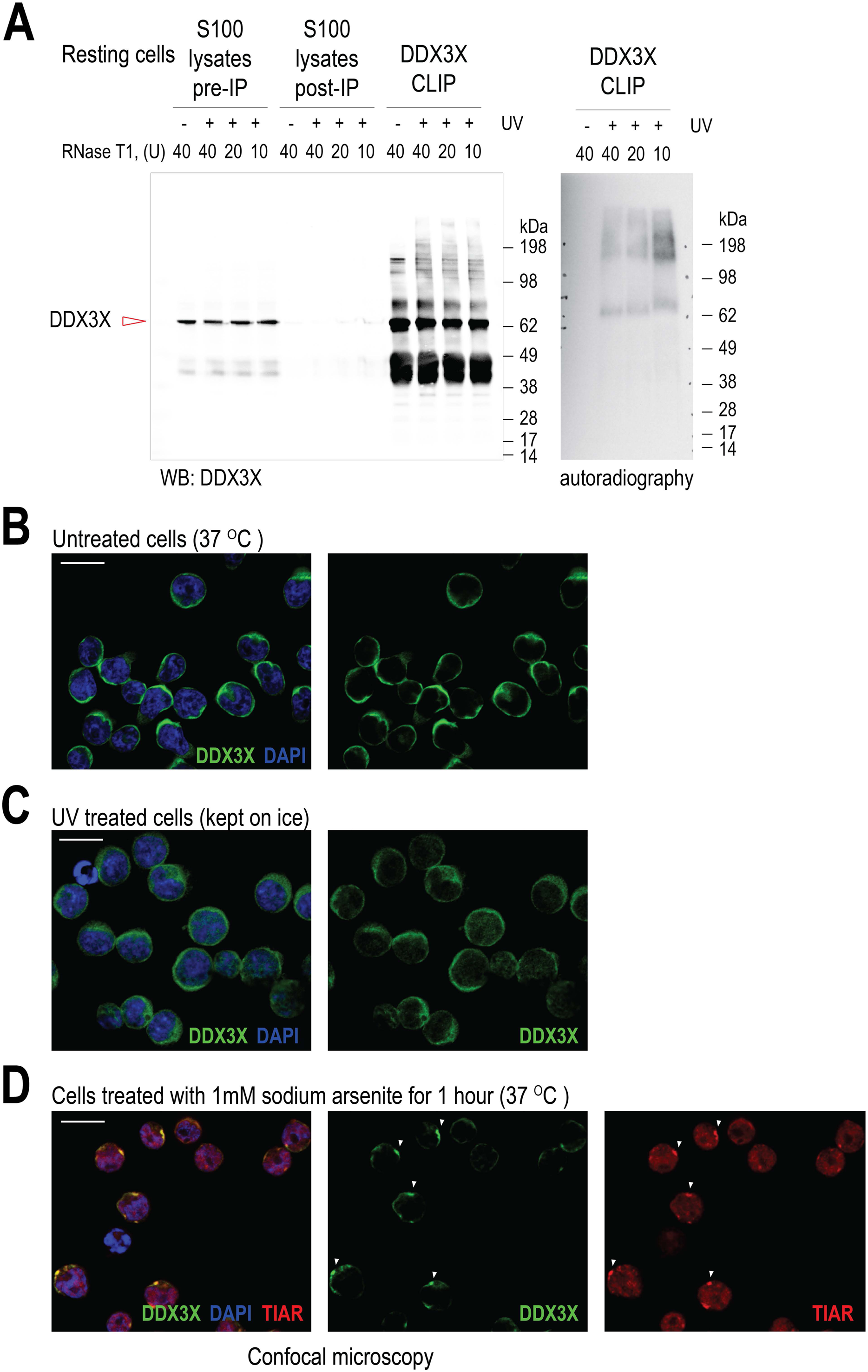
DDX3X CLIP optimization; immunofluorescence of untreated and UV treated 8E5 cells. (A) DDX3X immunoprecipitation, western blot and autoradiography, for DDX3X CLIP. Optimization of RNase T1 treatment of the cell lysate prior to immunoprecipitation. No UV treated cells are used as negative control. (B) DAPI and DDX3X immunofluorescence of untreated 8E5 cells grown at standard cell culture conditions. Scale bars: 15 μm (C) DAPI and DDX3X immunofluorescence of 8E5 cells after 3 exposures to UV (each at 400 mJ/cm^2^), with 30 sec intervals for cooling. Cells are kept on ice during this procedure, which lasts approximately 6 min in total. Scale bars: 15 μm (D) DAPI, DDX3X and TIAR immunofluorescence of 8E5 cells after treatment with 100 μM sodium arsenite for 1h. Note stress granule formation, in which DDX3X and TIAR colocalize (white arrowheads). Scale bars: 15 μm

**Figure S2.**
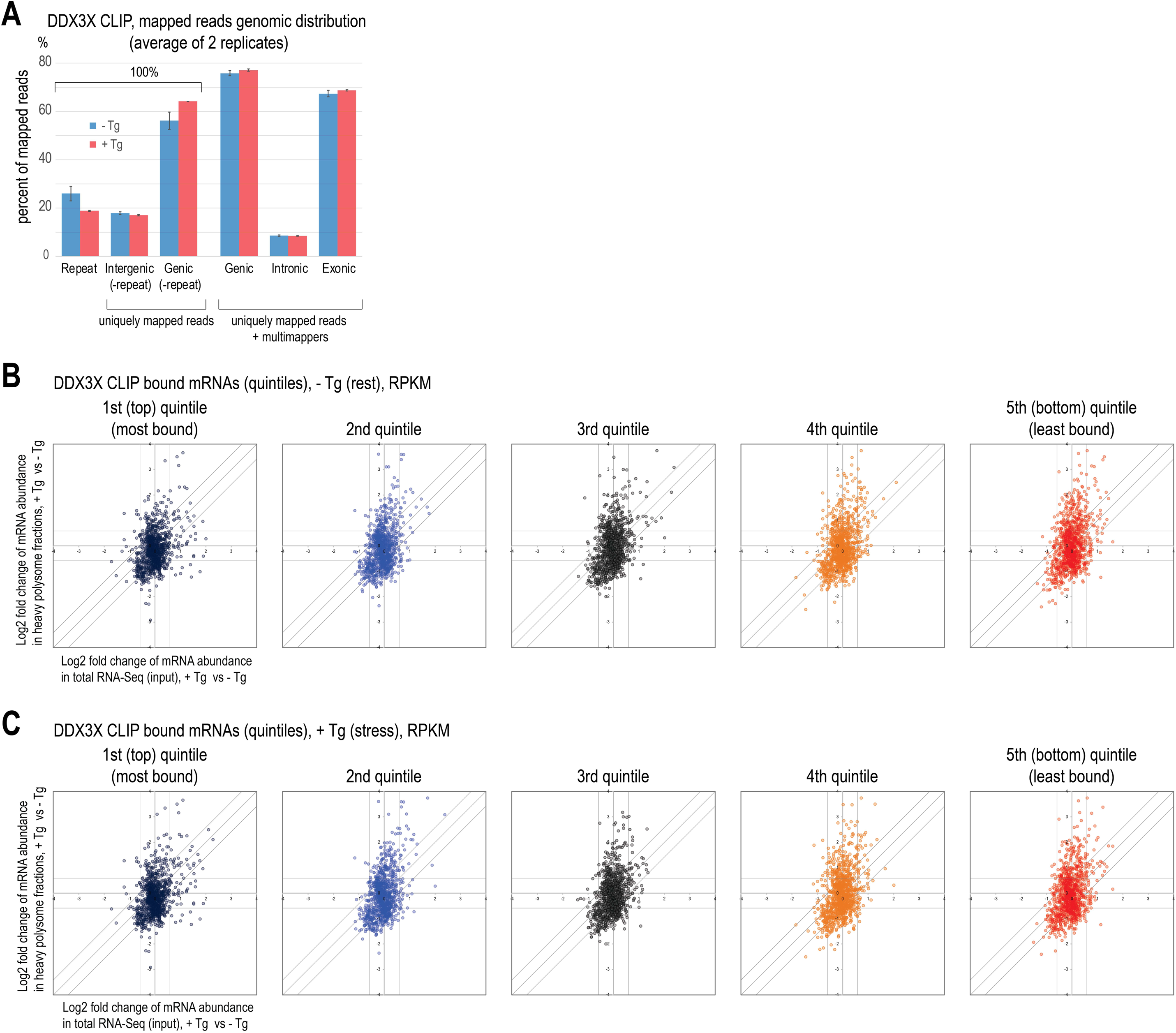
Polysome profiling scatterplots of mRNAs bound by DDX3X in MEFs, without or with Tg treatment. (A) Genomic distribution of DDX3X CLIP-Seq libraries. N=2, error bars: min-max. (B) mRNAs that are bound by DDX3X (CLIP-Seq abundance ≥ 1 RPKM) were ranked from highest to lowest CLIP-Seq abundance in-Tg treated cells, and split into five equal groups (quintiles). The mRNAs in these five separate groups were plotted as a scatterplot of polysome profiling in 1h Tg treated MEFs vs 0h Tg control MEFs, left to right most to least DDX3X bound mRNAs in-Tg cells, to explore the consequence of different levels of DDX3X binding of mRNAs in-Tg cells, upon Tg-induced ER stress. We observed that mRNAs from all seven categories of regulation are present in the polysome profile scatterplots of all quintiles, contradicting the hypothesis that the degree of DDX3X association (or dissociation) with an mRNA can predict how the mRNA translational regulation. (C) Same as in (A), but the DDX3X-bound mRNAs are split in quintiles based on their abundance in +Tg treated MEFs.

**Figure S3.**
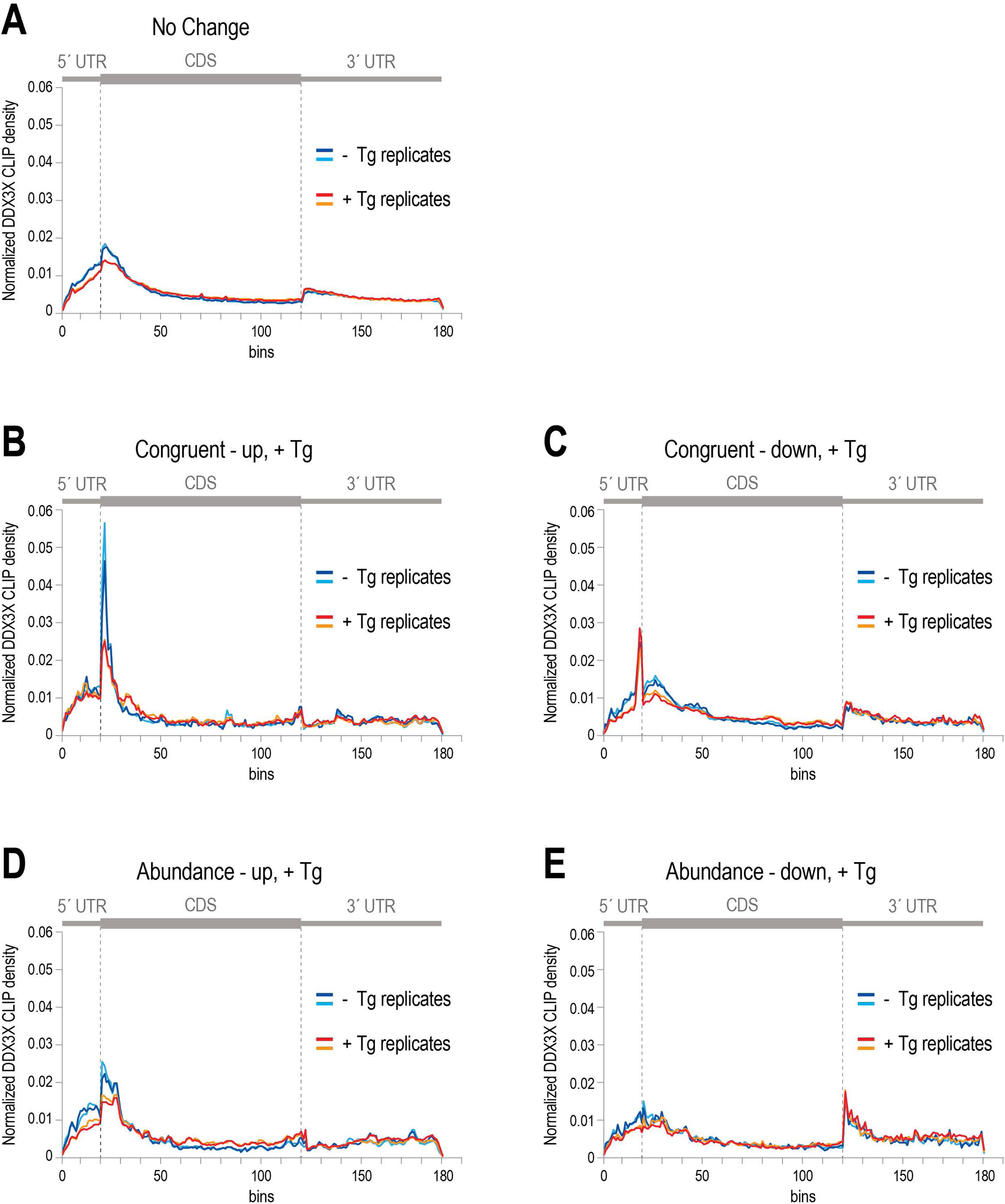
DDX3X CLIP-Seq in MEFs without and with Thapsigargin treatment (A) DDX3X CLIP density (CLIP in-Tg and +Tg treated cells in blue and red lines respectively) across the meta mRNA for mRNAs whose abundance and polysome association does not change in +Tg treated cells vs-Tg. (B) DDX3X CLIP density (CLIP in-Tg and +Tg treated cells in blue and red lines respectively) across the meta mRNA for mRNAs that are showing congruent increase of abundance and translation in +Tg treated cells vs-Tg. (C) DDX3X CLIP density (CLIP in-Tg and +Tg treated cells in blue and red lines respectively) across the meta mRNA for mRNAs that are showing congruent decrease of abundance and translation in +Tg treated cells vs-Tg. (D) DDX3X CLIP density (CLIP in-Tg and +Tg treated cells in blue and red lines respectively) across the meta mRNA for mRNAs that are showing increase of abundance in +Tg treated cells vs-Tg. (E) DDX3X CLIP density (CLIP in-Tg and +Tg treated cells in blue and red lines respectively) across the meta mRNA for mRNAs that are showing decrease of abundance in +Tg treated cells vs-Tg.

**Figure S4.**
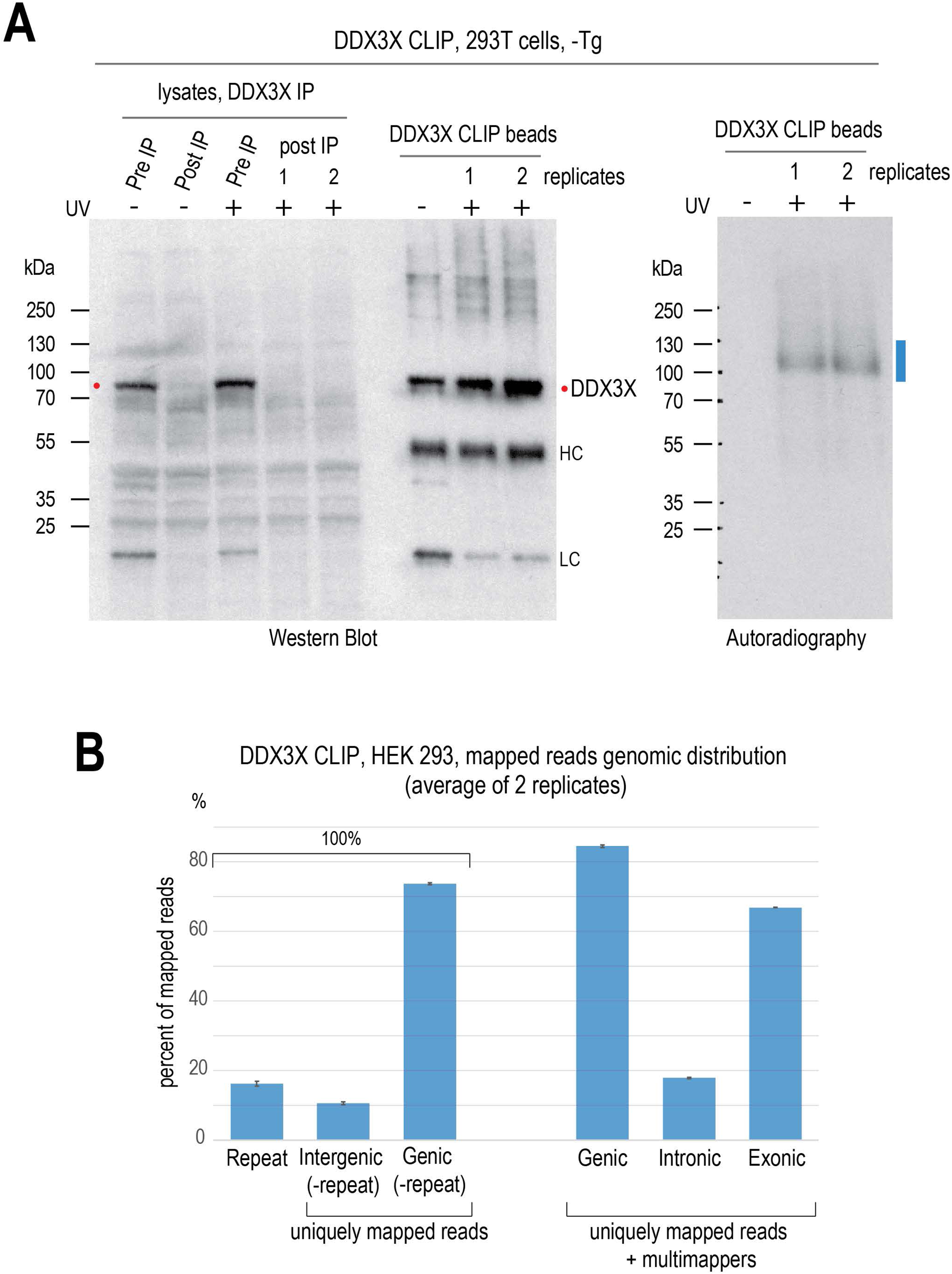
DDX3X CLIP-Seq in human 293T cells (A) DDX3X CLIP in 293T cells, not treated with Tg, two biological replicates. As a negative control DDX3X IP was performed in cells that were not exposed to UV crosslinking, resulting in complete absence of a radioactive signal in the autoradiography. DDX3X western blot signal is marked with a red dot, and crosslinked and labeled RNA-protein complexes extracted for CLIP-Seq library generation are marked with a blue line. (B) Genomic distribution of mapped reads of DDX3X CLIP-Seq libraries from control 0h Tg 293T cells. In the left part of the bar plot three mutually exclusive categories of genomic elements are shown, that account for all the mapped reads. The category “Repeat” contains reads mapped in multiple locations, including intergenic and genic sequences. In the right part of the plot, the percentage of the CLIP-Seq libraries mapped on annotated genes, introns and exons is shown. These include uniquely mapped and multimapping reads mapping exclusively in genes. Reads mapped in exons constitute the vast majority of DDX3X CLIP-Seq libraries. Average percentages of two replicate libraries per condition are shown; error bars are min and max values of the two replicates.

**Figure S5.**
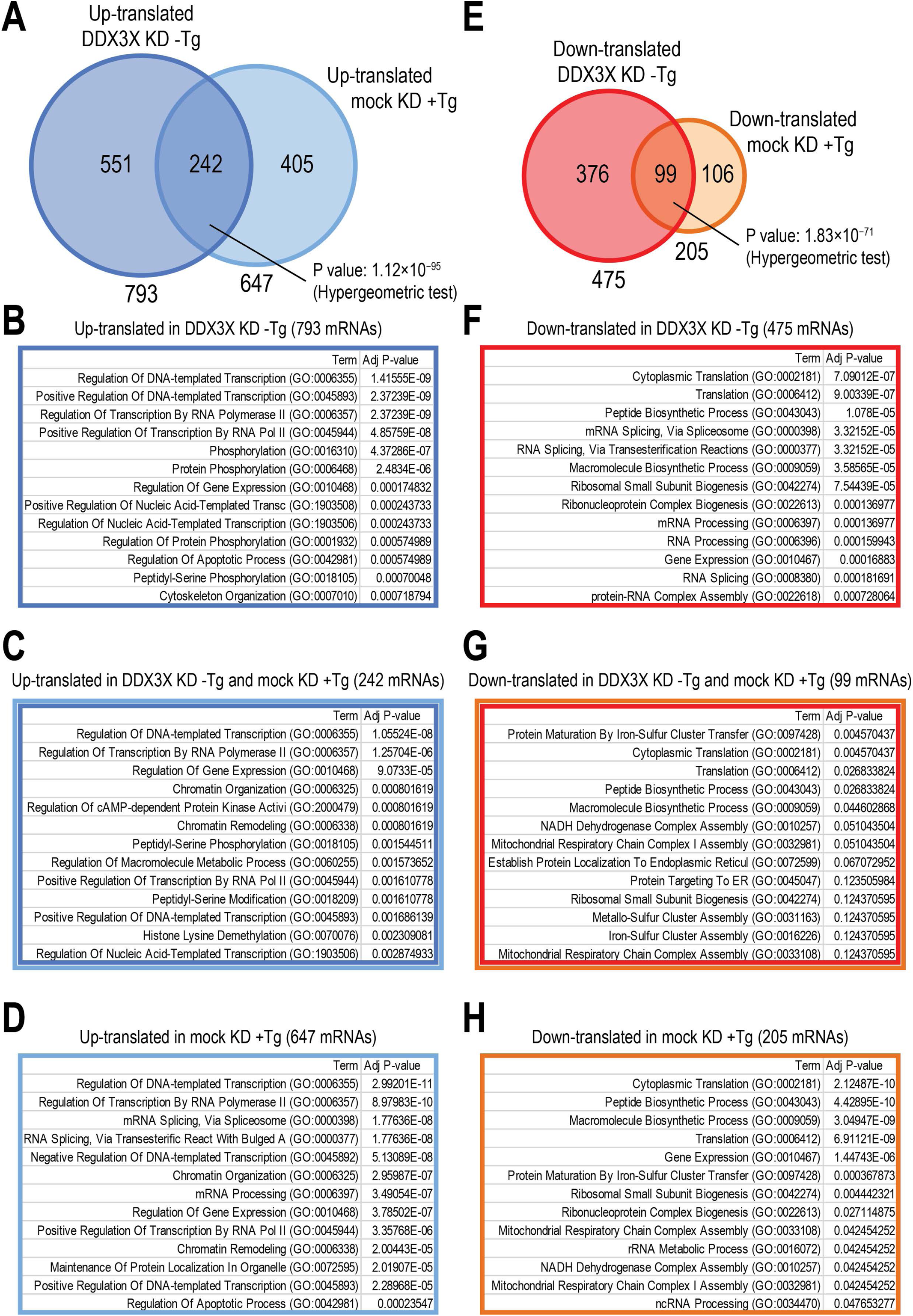
Homodirectional mRNA translational changes in DDX3X KD-Tg and mock KD +Tg suggests a role for DDX3X in suppressing stress-related mRNAs in non-stressed cells, and stimulating translation of mRNAs of “resting” character that are suppressed in stressed cells. A full list of enriched gene ontology categories for all gene lists in Figure S5 can be found in Table S3. (A) Venn diagram of up-translated mRNAs in DDX3X KD-Tg treated cells and in mock KD +Tg treated cells, and statistic test for the overlap as indicated. (B) Gene ontology analysis for the 793 up-translated mRNAs in DDX3X KD-Tg cells. Top 13 categories are shown, ranked by their p-value. (C) Gene ontology analysis for the 242 mRNAs that are up-translated in both DDX3X KD-Tg cells and in mock KD +Tg. Top 13 categories are shown, ranked by their p-value. (D) Gene ontology analysis for the 647 up-translated mRNAs in mock KD +Tg. Top 13 categories are shown, ranked by their p-value. (E) Venn diagram of down-translated mRNAs in DDX3X KD-Tg treated cells and in mock KD +Tg treated cells, and statistic test for the overlap as indicated. (F) Gene ontology analysis for the 475 down-translated mRNAs in DDX3X KD-Tg cells. Top 13 categories are shown, ranked by their p-value. (G) Gene ontology analysis for the 99 mRNAs that are down-translated in both DDX3X KD - Tg cells and in mock KD +Tg. Top 13 categories are shown, ranked by their p-value. (H) Gene ontology analysis for the 205 down-translated mRNAs in mock KD +Tg. Top 13 categories are shown, ranked by their p-value.

**Figure S6.**
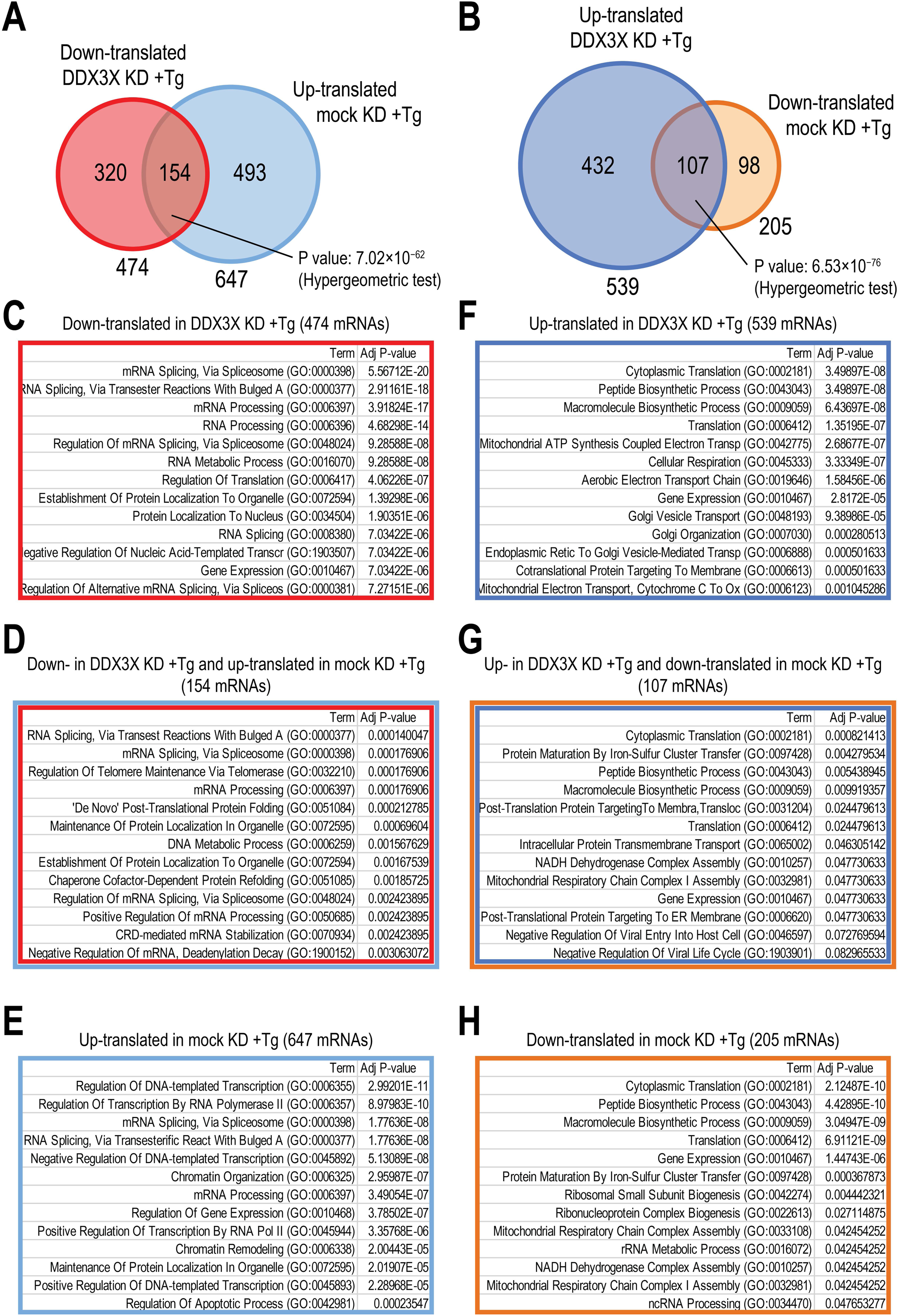
Heterodirectional mRNA translational changes in DDX3X KD +Tg and mock KD +Tg suggests a role for DDX3X in activating stress-related mRNAs in cells experiencing acute ER stress, and suppressing mRNAs of “resting” character. A full list of enriched gene ontology categories for all gene lists in Figure S5 can be found in Table S3. (A) Venn diagram of down-translated mRNAs in DDX3X KD +Tg treated cells and up-translated in mock KD +Tg treated cells, and statistic test for the overlap as indicated. (B) Venn diagram of up-translated mRNAs in DDX3X KD +Tg treated cells and down-translated in mock KD +Tg treated cells, and statistic test for the overlap as indicated. (C) Gene ontology analysis for the 474 down-translated mRNAs in DDX3X KD +Tg cells. Top 13 categories are shown, ranked by their p-value. (D) Gene ontology analysis for the 154 mRNAs that are down-translated in DDX3X KD +Tg cells and up-translated in mock KD +Tg. Top 13 categories are shown, ranked by their p-value. (E) Gene ontology analysis for the 647 up-translated mRNAs in mock KD +Tg. Top 13 categories are shown, ranked by their p-value. (F) Gene ontology analysis for the 539 up-translated mRNAs in DDX3X KD +Tg cells. Top 13 categories are shown, ranked by their p-value. (G) Gene ontology analysis for the 107 mRNAs that are up-translated in DDX3X KD +Tg cells and down-translated in mock KD +Tg. Top 13 categories are shown, ranked by their p-value. (H) Gene ontology analysis for the 205 down-translated mRNAs in mock KD +Tg. Top 13 categories are shown, ranked by their p-value.

**Figure S7.**
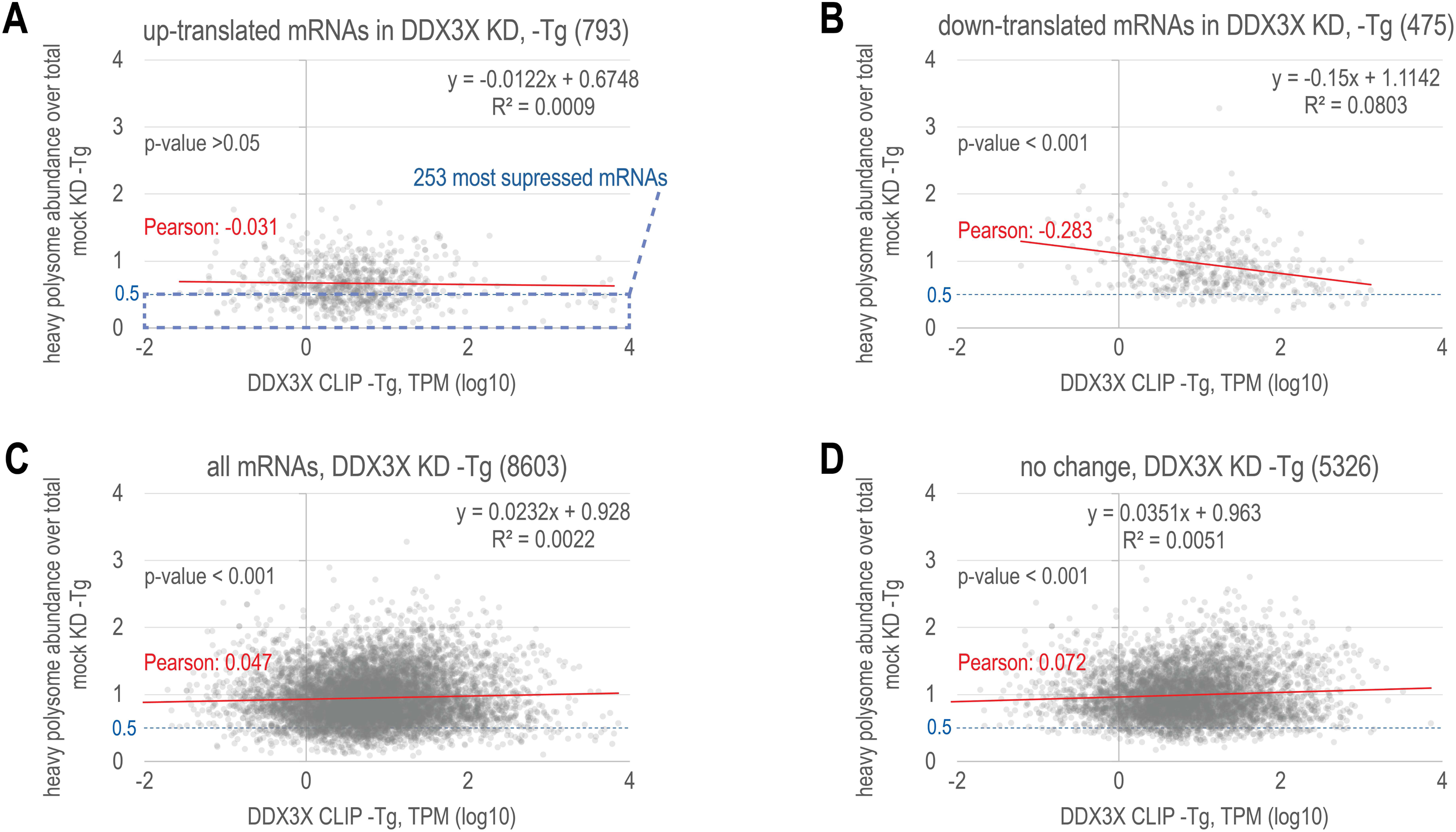
DDX3X CLIP-Seq abundance of an mRNA does not predict mode and degree of its translational regulation. (A) Scatterplot of DDX3X CLIP abundance (x-axis, average of two replicates, TPM) vs the ratio of heavy polysome over total RNA input abundances from the polysome profile of mock KD-Tg cells, for the 793 mRNAs showing increased polysome association (up-translation) in DDX3X KD-Tg (vs mock KD-Tg). The dashed box highlights the 253 most suppressed mRNAs (heavy polysomes over total abundance >0.5 that by definition in this category are up-translated in DDX3X KD-Tg. Pearson correlation (r) and trendline are shown in red, R^2^, trendline equation and p-value in grey. (B) Same as in A, but for the 475 mRNAs showing decreased polysome association (down-translation) in DDX3X KD-Tg (vs mock KD-Tg). (C) Same as in A, but for the 5326 mRNAs not showing any change in abundance or polysome association in DDX3X KD-Tg (vs mock KD-Tg).

**Figure S8.**
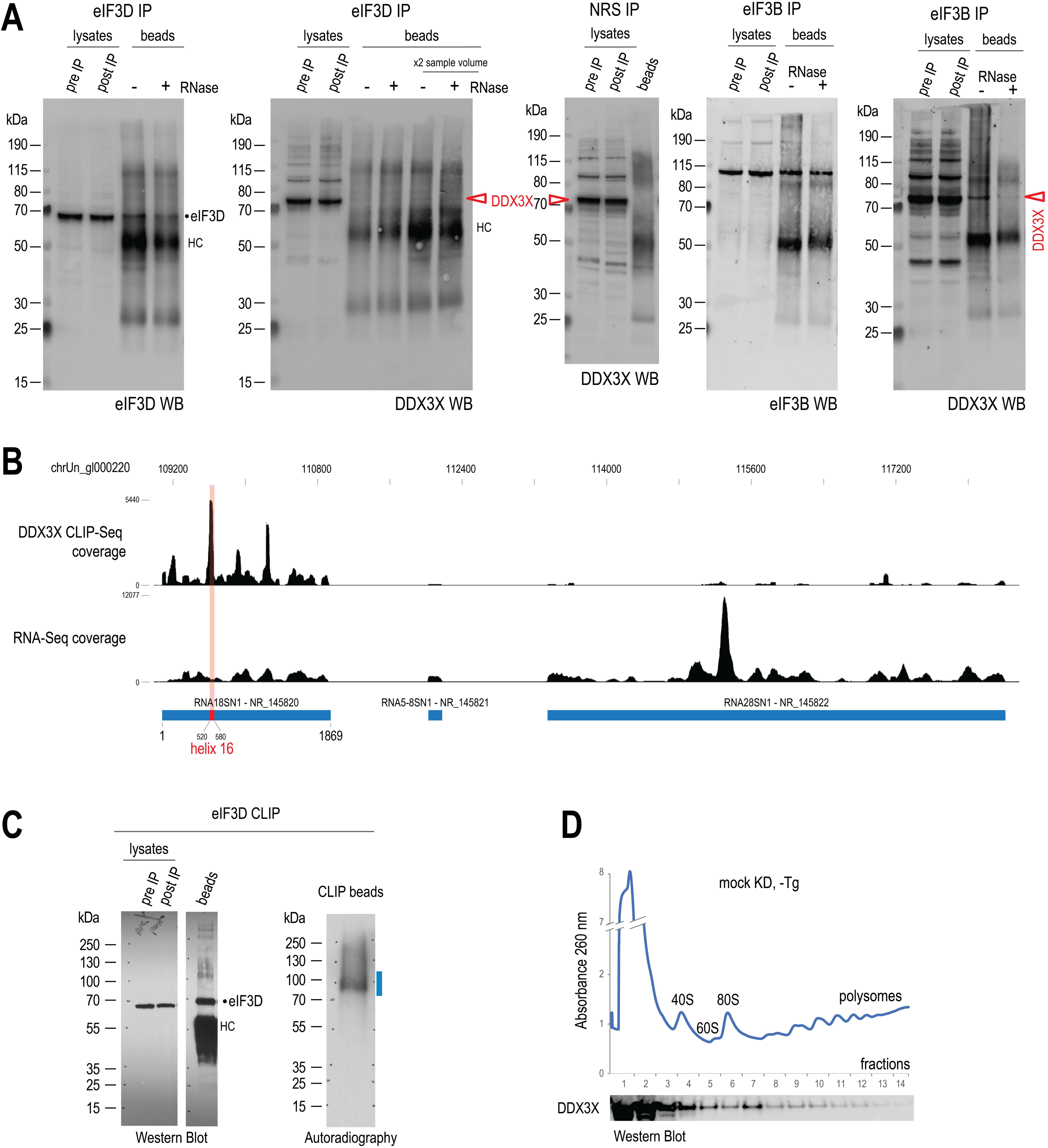
Interactions of DDX3X with the translation machinery (A) Co-immunoprecipitation of DDX3X with eIF3D and eIF3B, without and with RNase T1 treatment of IP beads before protein elution. NRS IP as a negative control. A red triangle marks the DDX3X band in beads eluate samples. (B) UCSC Genome browser snapshot of human 45S rRNA locus (hg19). Top track: DDX3X CLIP-Seq coverage; bottom track, RNA-Seq coverage. Genomic coordinates are shown at the top, ribosomal RNAs and their designations are shown at the bottom of the tracks. The location of helix 16 on 18S rRNA is marked in red. We note the preferential binding of DDX3X on 18S rRNA rather than 28S and 5.8S rRNA, and the prominent peak on Helix 16, also when compared to RNA-Seq coverage. (C) eIF3D CLIP, in 293T-Tg cells; WB of the lysates and the IP beads (left), and autoradiography (right) of RNA-protein complexes, with the membrane area from which RNA was extracted highlighted in blue. (D) DDX3X WB with sucrose gradient fractions from mock KD-Tg cells, showing presence of DDX3X in polysome fractions.

**Figure S9.**
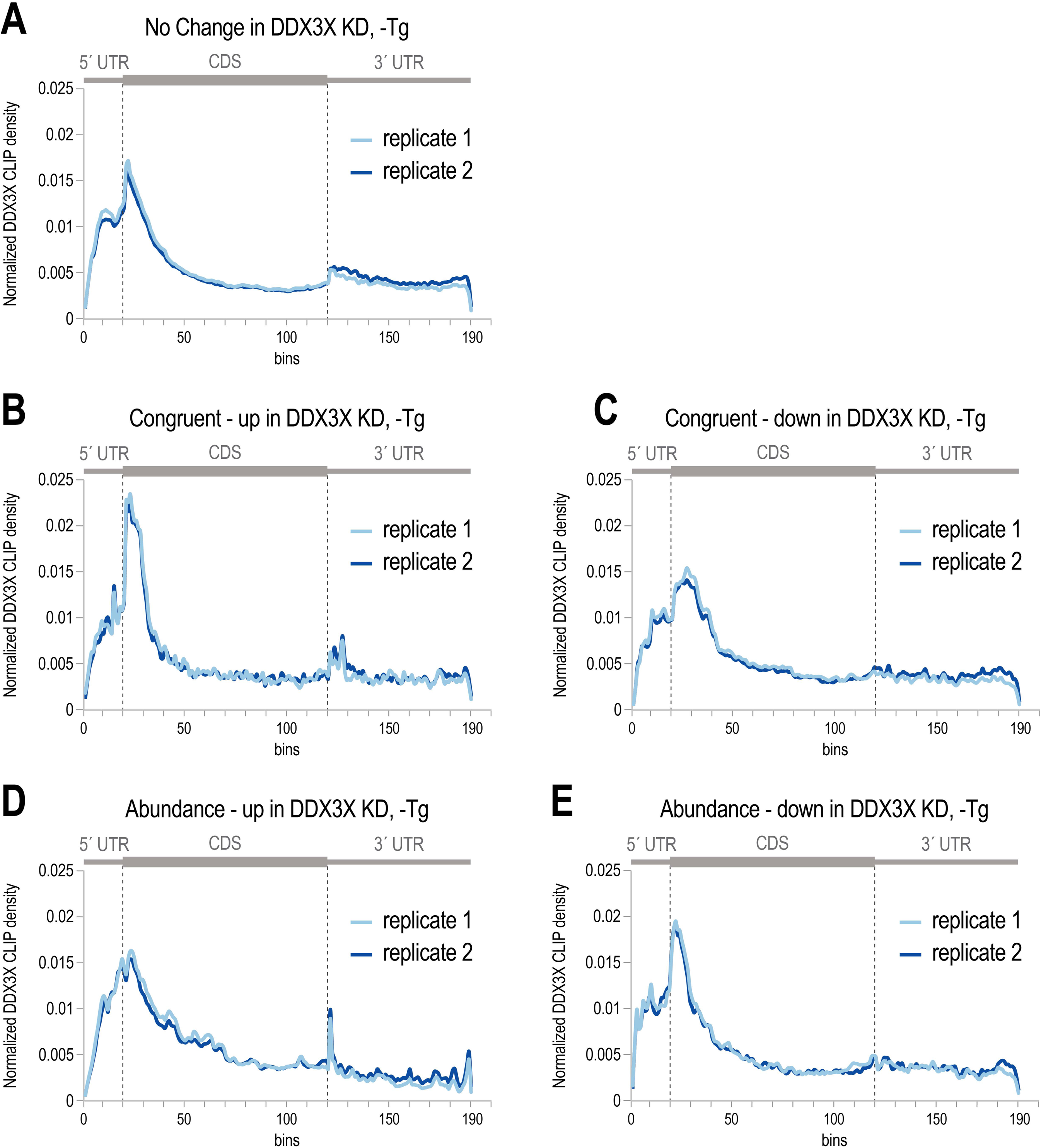
DDX3X CLIP-Seq 293T cells, meta mRNA plots (A) DDX3X CLIP density (two biological replicates,-Tg treated cells) across the meta mRNA for mRNAs whose abundance and polysome association does not change upon DDX3X KD. (B) DDX3X CLIP density (two biological replicates,-Tg treated cells) across the meta mRNA for mRNAs whose abundance and polysome association increases in a congruent manner upon DDX3X KD. (C) DDX3X CLIP density (two biological replicates,-Tg treated cells) across the meta mRNA for mRNAs with congruent decrease of abundance and polysome association upon DDX3X KD. (D) DDX3X CLIP density (two biological replicates,-Tg treated cells) across the meta mRNA for mRNAs whose abundance increases upon DDX3X KD. (E) DDX3X CLIP density (two biological replicates,-Tg treated cells) across the meta mRNA for mRNAs whose abundance decreases upon DDX3X KD.

**Table S1.** Number of reads for the deep-sequencing libraries generated in this work.

**Table S2.** mRNA transcript TPM values, and fold changes of total RNA and heavy polysome associated mRNA abundances for the polysome profiling experiments performed in this work. The regulatory consequence for each mRNA transcript in the five different experiments is also shown. **Table S3.** Lists of up-and down-translated mRNAs for the indicated experiments, and Gene Ontology analysis (Enricher).

**Table S4.** List of proteins and numbers of identified peptides for each protein from DDX3X immunoprecipitation and antigenic peptide elution.

## Notes

### Competing Interest Statement

The authors have declared no competing interest.

